# High-throughput mapping of 6,888 *RAD51D* variants identifies distinct biochemical functions needed for homologous recombination and olaparib response

**DOI:** 10.64898/2026.01.11.698865

**Authors:** Kristie E. Darrah, Shelby L. Hemker, Yashpal Rawal, Noah J. Goff, Phoebe Parker, Gayatri Ganesan, Caleb M. Stratton, Katherine Oppenheimer, Ella Roberts, Elena Glick, Nicole Banks, Arjun Kumar, Silvia Casadei, Matthew W. Snyder, Katherine Nathanson, Susan M. Domchek, Lea M. Starita, Shaun K. Olsen, Patrick Sung, Jacob O. Kitzman, Kara A. Bernstein

**Affiliations:** Department of Biochemistry and Biophysics, University of Pennsylvania Perelman School of Medicine, Philadelphia, PA, USA; Department of Human Genetics, University of Michigan Medical School, Ann Arbor, MI 48109, USA; Gilbert S. Omenn Department of Computational Medicine & Bioinformatics, University of Michigan Medical School, Ann Arbor, MI 48109, USA; Department of Biochemistry and Structural Biology and Greehey Children’s Cancer Research Institute, University of Texas Health Science Center at San Antonio, San Antonio, TX, USA; Department of Pharmacology and Chemical Biology, University of Pittsburgh, School of Medicine, Pittsburgh, PA, USA; Department of Medicine, Basser Center for BRCA, University of Pennsylvania Perelman School of Medicine, Philadelphia, PA 19104, USA; Department of Genome Sciences, University of Washington, Seattle, WA 98195, USA; Brotman Baty Institute, Seattle, WA 98195, USA

**Keywords:** Variants of Unknown Significance, Multiplex Assay of Variant Effect, Homologous Recombination, RAD51D, RAD51, RAD51 Paralog, BCDX2, X3CDX2, XRCC3 complex

## Abstract

The tumor suppressor *RAD51D* is essential for homologous recombination (HR). Pathogenic variants in *RAD51D* are associated with breast and ovarian cancers. However, most clinical missense variants are of unknown significance. We performed a multiplex assay of variant effect to test 6,888 *RAD51D* coding variants for loss-of-function. The resulting variant-to-function map perfectly separates known pathogenic and benign variants and is validated by orthogonal HR and biochemical assays across 70 clinical variants. Our screen shows that variants in the DNA-binding or ATPase core most severely compromise HR, and we identify the RAD51D-RAD51C interface within the BCDX2 complex as essential for regulating its ATPase activity. We hypothesize that, paradoxically, the primary function of RAD51D is to slow the ATPase activity of BCDX2, thereby allowing sufficient time and space for RAD51 filament assembly. Together, we identify hotspots of deleterious *RAD51D* variants and uncover the mechanisms by which variants compromise its biochemical functions.

**Highlights:**

- Used a multiplexed assay of functional effect (MAVE) to assess the functionality via olaparib sensitivity of 6,888 *RAD51D* coding variants, which can be used for variant classification
- Provided cellular functional analysis for 70 clinically-identified breast and ovarian cancer *RAD51D* variants
- Identified key regions and enzymatic activities of RAD51D critical for its function in the BCDX2 and the X3CDX2 complexes
- Determined mechanism of RAD51D-mediated regulation of BCDX2 ATPase activity

## Introduction

Defects in homologous recombination (HR) genes are closely associated with breast and ovarian cancers, as well as Fanconi anemia, a rare disease characterized by cancer and bone marrow failure^1^. As a result, numerous HR genes are routinely included on cancer screening panels^2^ and are considered actionable. Among these, *RAD51D* is a paralog of the recombinase *RAD51,* and plays an essential role in the repair of DNA double-strand breaks through the HR pathway and in the repair of replicative damage. Loss of *RAD51D* function often occurs through germline inheritance of a single deleterious variant, with the eventual somatic loss of the second wild-type copy of *RAD51D* contributing to tumor development^3,4^.

Although disruption of the HR pathway is well-known as a cancer risk factor, defining which variants in HR genes are pathogenic poses a significant barrier for patients and related family members, who may share risk variants. Family history of *RAD51D*-associated cancers is often challenging to recognize, given the lower penetrance compared with other hereditary breast and ovarian cancer genes such as *BRCA1* and *BRCA2*^5^. While nonsense variants can often be classified as pathogenic, missense variants are routinely identified in the clinic and most remain as variants of unknown significance (VUS). In *RAD51D* alone, >850 missense variants are reported in ClinVar, with >96% having VUS or conflicting interpretations (**Figure 1a**). The absence of functional characterization for these VUS variants poses a significant barrier in clinical decision-making, hindering the ability to identify which patients require increased surveillance and/or specialized treatment strategies.

**Figure 1:**
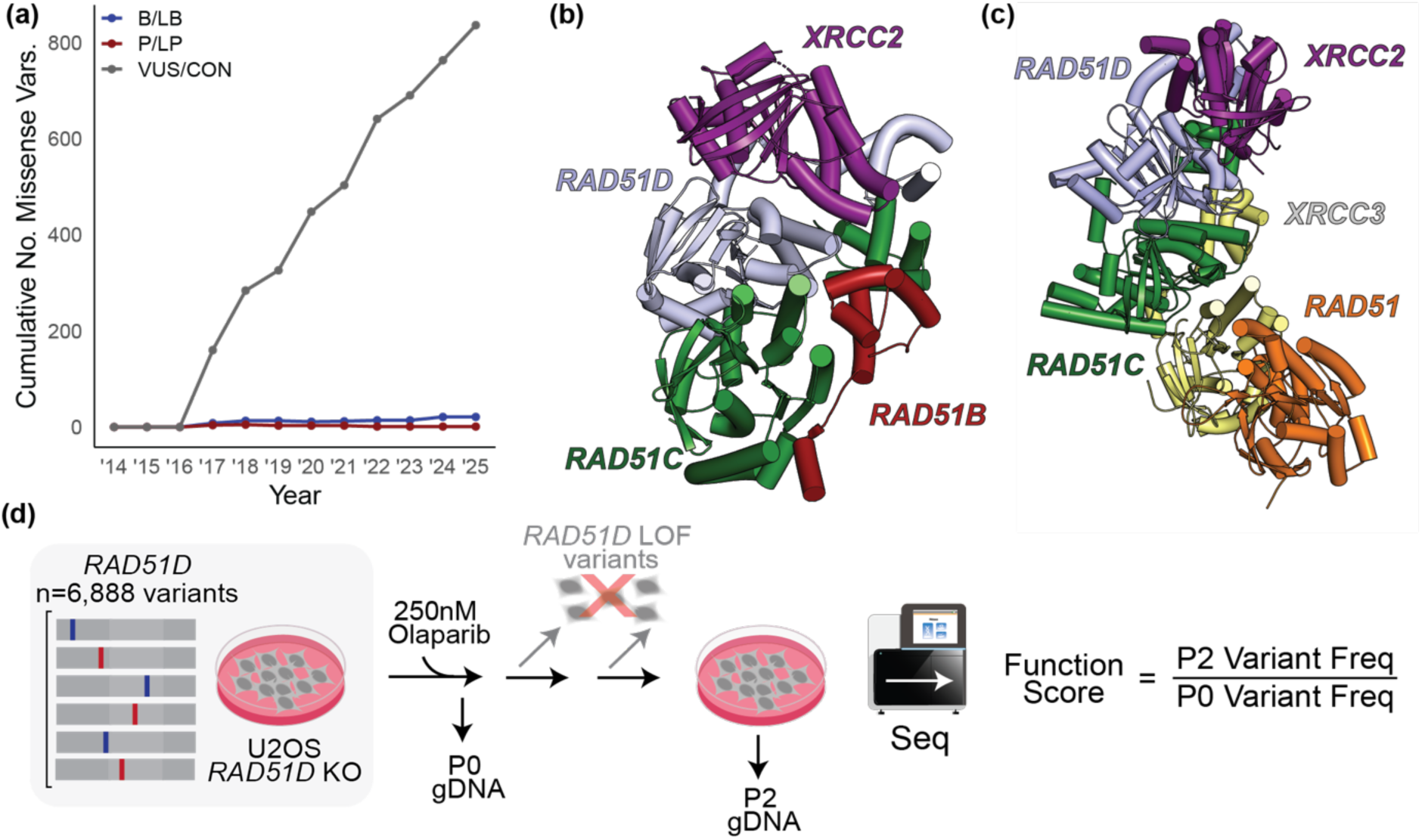
Overview of pooled variant-to-function mapping in RAD51D. **(a)** ClinVar classifications of *RAD51D* missense variants each year. P/LP = Pathogenic/Likely Pathogenic, B/LB = Benign/Likely Benign, VUS/CON = unknown significance/conflicting reports. (**b-c**) RAD51D forms an obligate heterodimer with XRCC2 to form the (**b**) BCDX2 (PDB: 8GBJ) or (**c**) XRCC3 (X3CDX2) complexes (PDB: 9SVX) through its interaction with RAD51C. (**d**) Schematic of pooled *RAD51D* variant function assay. *RAD51D* mutant library cell population is treated with 250 nM olaparib over several passages, selecting for cells with normal *RAD51D* function. *RAD51D* variants are quantified within the starting (P0) and final (P2) cell populations by sequencing and scored by their relative enrichment in P2 versus P0.

Multiplex assays of variant effect (MAVE) have recently enabled high-throughput measurement of variant functional consequences across clinically relevant and actionable human disease-associated genes^6,7^. Following genome editing or stable expression of one variant per cell, a functional selection is applied based upon cell fitness or resistance to genotoxic drugs, after which deep sequencing identifies variants that become enriched or depleted. This approach has been successfully used to identify functionally disruptive variants in HR genes including *BRCA2,* and the *RAD51* paralog *RAD51C*^8–10^.

Central to understanding the clinical impact of *RAD51D* is understanding its HR function. RAD51D forms a heterotetrameric complex with the RAD51 paralogs RAD51B, RAD51C, and XRCC2 (herein BCDX2; **Figure 1b**). Additionally, RAD51D is a key member of another RAD51 paralog-containing complex, the XRCC3 complex (herein X3CDX2), which comprises XRCC3, RAD51C, RAD51D, and XRCC2 (^11^; **Figure 1c**). In ssDNA-bound structures of both BCDX2 and X3CDX2 complexes, the DNA-binding loops L1 and L2 of RAD51D play a critical role, as mutations in these regions impair BCDX2’s ability to bind ssDNA^11,12^. Although all of the paralogs share 20-30% structural similarity to RAD51^13,14^, particularly in conserved ATPase core domains and motifs, only the RAD51B-RAD51C (herein BC) heterodimer hydrolyzes ATP, whereas the RAD51D-XRCC2 (herein DX2) heterodimer regulates BC ATPase activity through an unknown mechanism^12^. Not surprisingly, previous functional analysis of the related RAD51C protein has identified the ATPase domains as central for its HR function, and deleterious variants in these regions have been identified in breast and/or ovarian cancers^15,16^. Although *RAD51C* variants have been identified in approximately 3% of HR deficient ovarian cancers, *RAD51D* is mutated more frequently, occurring in 5% of such cases^17^. In contrast to *BRCA2* and *RAD51C*, only a handful of variants in *RAD51D* have had any prior functional analysis to date, and the impact of clinically identified breast and ovarian cancer variants on HR is largely unknown (summarized in **Supplementary Table 1**). Furthermore, the functional regions and domains essential for RAD51D activity, and the manner in which RAD51D, together with XRCC2, regulates BC ATPase activity, have yet to be elucidated.

Here, we used a MAVE to analyze 6,888 single-codon *RAD51D* variants for loss of viability using sensitivity to the poly (ADP-ribose) polymerase inhibitor (PARPi), olaparib, which is currently used to treat HR-deficient ovarian cancers^18^. The resulting variant-to-function map provides measurements for 97% of all single-residue variants, offering a global view of missense constraint concentrated in the ATPase-like and DNA-binding domains. Next, 70 variants previously identified in breast and/or ovarian cancers were selected for subsequent functional studies of HR activity and complex interactions involving BCDX2 and X3CDX2. Having found that the results of the MAVE analysis strongly correlated with individual variant mechanistic studies, we identified seven residues in RAD51D critical for HR function and performed biochemical analyses to investigate their unique mechanistic contributions to BCDX2 complex formation, DNA binding, and ATPase regulation. We found that six of the seven deleterious *RAD51D* cancer-identified variants alter the regulation of BC ATPase activity in the BCDX2 complex, either by attenuating RAD51D interaction with RAD51C or by reducing BCDX2 DNA binding affinity. Lastly, we find that six of these variants impact the formation of the X3CDX2 complex. Together, our work is informative in identifying and reclassifying deleterious variants in *RAD51D*, providing mechanistic insight into how the disruption of these functional hotspots contributes to RAD51D function as part of the BCDX2 and X3CDX2 complexes.

## Results

### A variant-to-function map of all possible *RAD51D* single-codon variants

Since *RAD51D* is essential in non-transformed cells, we used the well-characterized osteosarcoma cell line U2OS in which *RAD51D* has been knocked out by CRISPR/Cas9 (herein U2OS *RAD51D* KO)^19^. It was previously shown that restoring RAD51D expression in these knockout cells rescues HR function and resistance to PARP inhibition through olaparib treatment^19,20^. We leveraged this assay to systematically test missense variants in the *RAD51D* gene (**Figure 1d**). Mutational libraries were stably integrated into U2OS *RAD51D* KO cells such that each resulting cell expressed a single *RAD51D* variant. We performed saturation mutagenesis of the *RAD51D* cDNA and obtained 97.6% coverage of 6,888 possible single-codon variants present at a frequency of ≥ 1/10,000 (**Supplementary Figure 1**). These mixed cellular populations were then treated with 250 nM olaparib for two passages to enrich for cells expressing functionally normal *RAD51D* mutants. Variant frequencies were quantified by deep amplicon sequencing before and after selection, and used to calculate a function score for each variant, with negative scores indicating variants that became depleted during olaparib treatment.

As expected, premature truncation variants showed strong depletion, with stop-gain variants prior to residue 305 having a mean score of −0.989. These were perfectly separated from the scores for synonymous variants (mean: 0.023; area under the precision-recall curve, prAUC=1.00). Notably, C-terminal truncations from residue 306 onward scored comparably to synonymous variants, consistent with this mostly unstructured region being dispensable in our assay. Variants were classified as neutral (NEU), intermediate (INT), or loss-of-function (LOF), using function score cutoffs set by the 2.5% and 97.5% percentiles of synonymous and nonsense variants, respectively (**Methods**). To be classified as INT (function scores between −0.170 and −0.446) or LOF (function score below −0.446), variants also had to pass a significance cutoff (local false sign rate, lfsr ≤0.01). As a result, we classified 74.5% of measured *RAD51D* missense variants (n=4,593) as NEU (mean score: −0.07), 16.8% as LOF (n=1,038; mean score: −0.78), and 8.6% as INT (n=536; mean score: −0.31) (**Figure 2a**). Supporting these classifications’ accuracy, all scored synonymous variants were classified as NEU, while nearly every nonsense variant prior to codon 305 (304/305, 99.7%) scored as INT or LOF.

**Figure 2:**
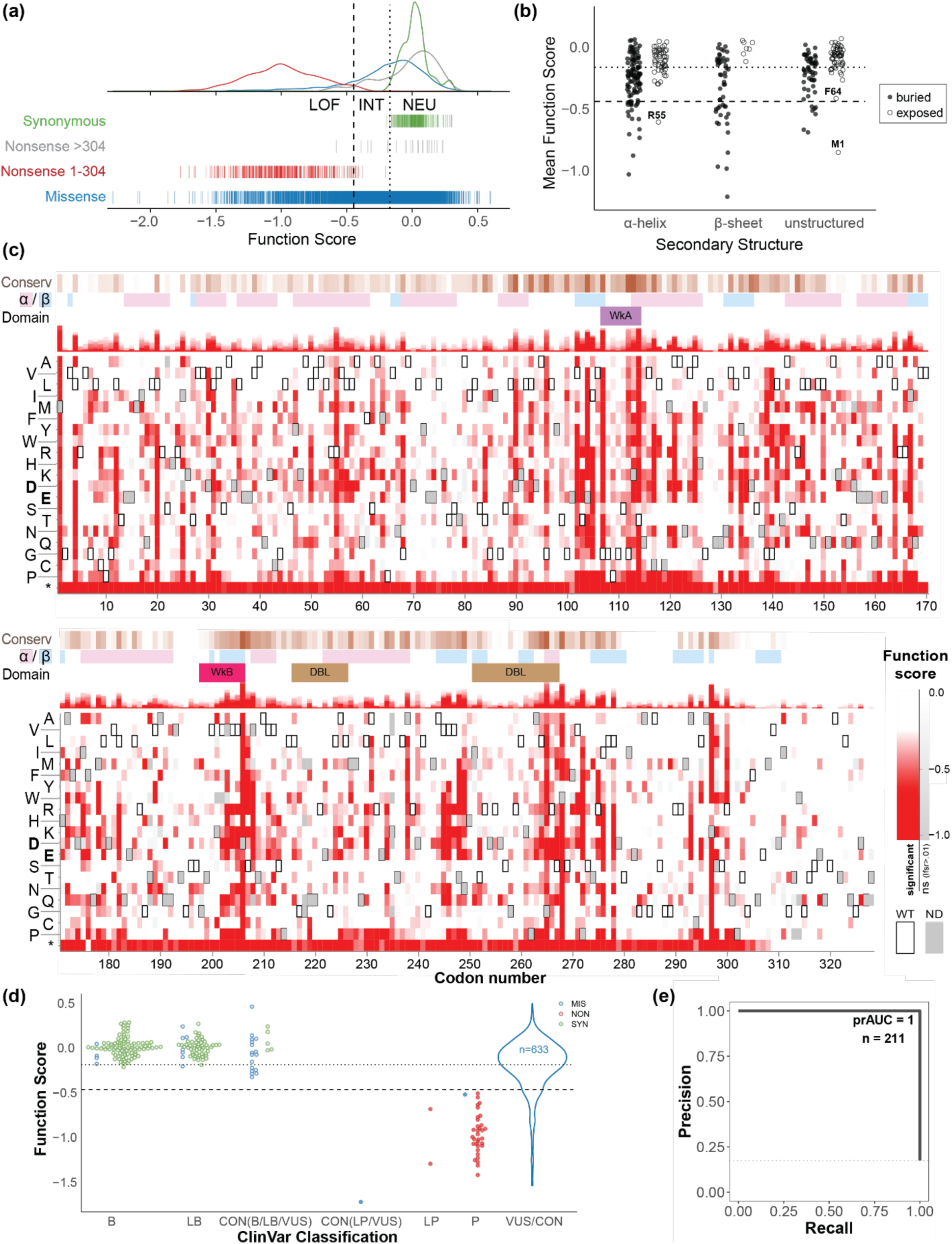
A variant-to-function heatmap of RAD51D. (**a**) Distributions of function scores by variant class; nonsense variants in codons 1–304 are perfectly separated from synonymous variants. Vertical lines denote cutoffs for function classifications (LOF, loss-of-function; INT, intermediate; NEU, neutral). (**b**) Per-residue mean missense function scores, grouped by residue surface area exposure status and secondary structure. Unique LOF exposed residues are labeled. (**c**) Variant-to-function heatmap across RAD51D, shaded by function score (white, WT-like; red, null-like) for each mutant amino acid (rows) at each codon position (columns). Variants that do not pass multiple testing correction (lfsr>0.01) are shaded from white to gray, WT residues are boxed, and dark gray denotes missing data. Tracks above heatmaps, from top to bottom: conservation score, protein secondary structure, key domains. Asp (D) or Glu (E) are shown in boldface to highlight stronger effects of these substitutions at some sites. (**d**) Function scores separate pathogenic from benign variants among single-residue variants reported in ClinVar, and provide evidence for missense VUS and those with conflicting reports (all variants plotted SpliceAI score <0.2). Point color denotes variant type, and filled/open points denote statistical significance (i.e. filled: lfsr ≤0.01). (**e**) Precision-recall curve showing classification performance between a set of 211 SNVs with confident P/LP and B/LB variant classification, included in (**a**).

Missense intolerance was enriched in structured, annotated domains of RAD51D (**Figures 2b-c, 3**). As expected, variants at buried residues were on average more deleterious than those at surface-exposed residues (mean function scores: −0.272 vs −0.113, Welch’s *t*: P<2.2×10^-16^; **Figure 2b**). However, we uncovered surface exposed residues near RAD51C or XRCC2 interfaces at which variants were highly deleterious (**Figure 3a,c,e**). Structural analysis identified several residues that make contact with DNA, ATP, or other proteins via hydrogen bonding or salt bridges that were largely intolerant to substitution (mean function score < −0.446, **Supplementary Figure 2b**). We found six highly constrained residues at which two or fewer variants were tolerated as NEU or INT. These appeared at well-conserved sites including the Walker A motif, where the conserved terminal threonine (T114) could be replaced only by serine, notable for being the other final amino acid of the classical Walker A motif^21^. Similarly, the Walker B motif ends in a conserved aspartic acid (D206) at which 18 of 19 substitutions exhibited strong LOF (mean function score: −1.22), except the chemically similar glutamic acid, which was less severe (score: −0.52). Residues near the ssDNA backbone (263-267) showed strong intolerance to Glu and Asp variants, which would be expected to repel the negatively charged backbone (**Figure 3f, Supplementary Figure 2a**). Glycine 265 was completely substitution-intolerant, suggesting it may provide functionally critical flexibility for fitting around the ssDNA backbone.

**Figure 3:**
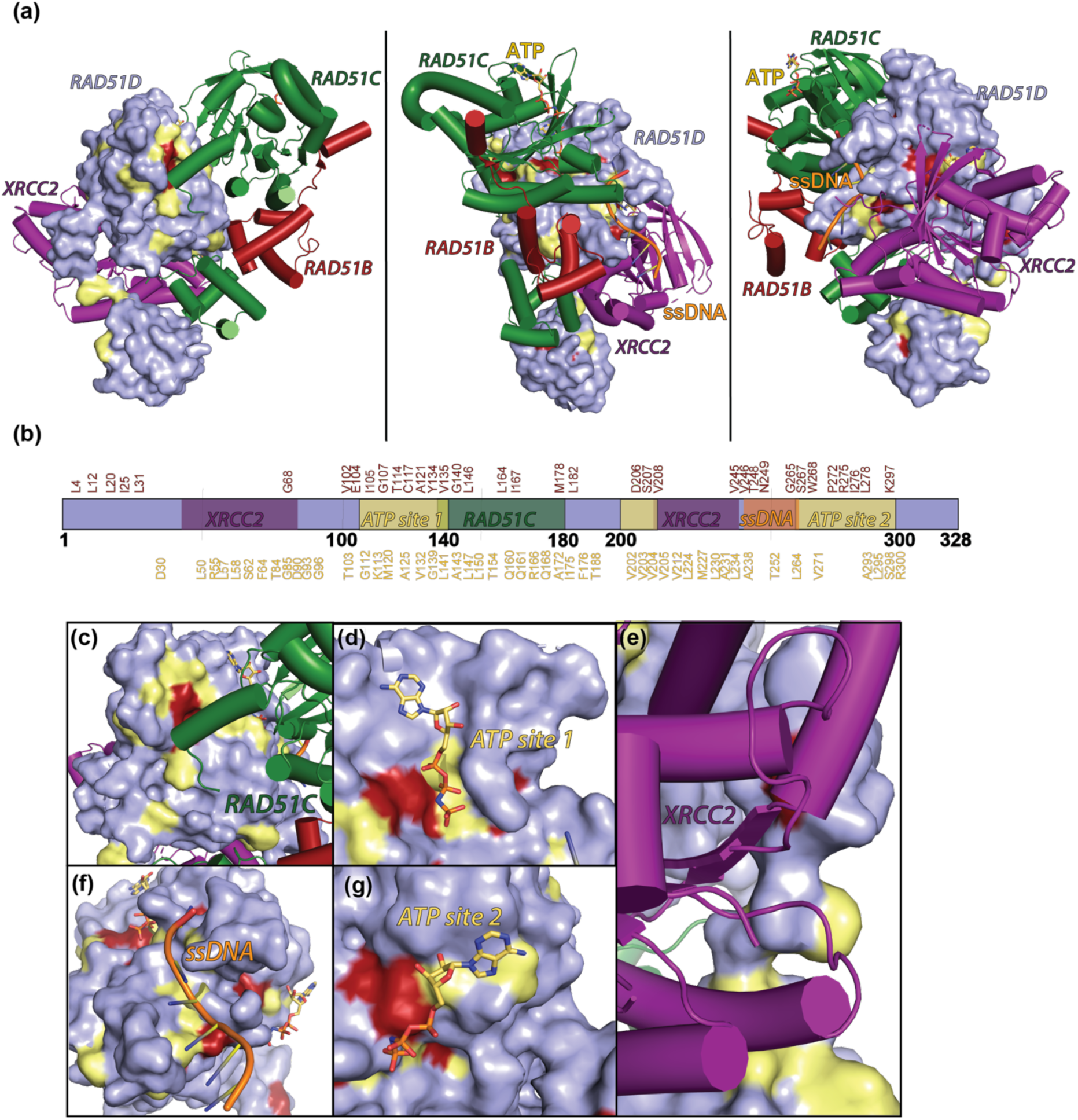
High-throughput screening reveals disrupted RAD51D interaction surfaces. (**a**) Surface representation of RAD51D within the BCDX2 complex (PDB: 8GBJ). Surface residues are colored according to either red (50% pathogenicity) or yellow (20% pathogenicity). Cartoon representations of XRCC2 (purple), RAD51D (light blue), RAD51C (green), and RAD51B (red) are shown consistently throughout. Bound ATP molecules are represented as yellow-orange sticks. (**b**) Specific loss-of-function variants (red, 50% pathogenicity) listed above or (yellow, 20% pathogenicity) listed below the RAD51D schematic are shown with the indicated interaction interfaces indicated. (**c-g**) Binding sites for RAD51C (**c**), ATP site 1 (**d**), XRCC2 (**e**), ssDNA (**f**), and ATP site 2 (**g**).

### MAVE scores provide strong evidence for *RAD51D* variants

To compare our function scores to existing clinical variant classifications, we curated a set of control variants from ClinVar. We identified 211 unique variants at the protein level, with B/LB (n=155) or P/LP (n=36) classification, along with an additional 28 “soft” conflicts meaning that there were two of more submissions with the same definitive classification (pathogenic or likely pathogenic (P/LP) or benign or likely benign (B/LB), along with VUS. To gain calibration power, we resolved these conflicts as P/LP (n=1) or B/LB (n=27; **Supplementary Table 5**). We observed perfect concordance with the clinical classifications; all 37 pathogenic or likely pathogenic (P/LP) variants scored as LOF and all 174 benign or likely benign (B/LB) variants scored as NEU in our MAVE (**Figure 2d-e**). The benign polymorphism R165Q scored as NEU, consistent with its high population allele frequency (gnomAD MAF=14.1%). Only two missense variants have been classified as pathogenic or likely pathogenic (P/LP and VUS in ClinVar), and both were correctly identified as LOF, with S207L having comparatively mild effects (score: −0.503) and S207W showing more severe impairment (score: −1.703). Additionally, we compared our function scores to AlphaMissense, a computational tool developed to predict missense variant pathogenicity from AlphaFold structural predictions and sequence conservation^22^. Notably, 35% of the variants called by AlphaMissense as pathogenic with high confidence (probability ≥0.9) were classified as functionally normal in our screen, suggesting a potentially high rate of false positives among these computational predictions (**Supplementary Figure 2c**).

To estimate the strength of evidence provided by our MAVE functional scores, we translated the functional scores into evidence for clinical variant classification following the OddsPath framework recommended by ClinGen^23^. OddsPath is a Bayesian odds ratio used to assign evidence based on the ability of the assay to correctly identify previously classified pathogenic as functionally abnormal and benign variants as functionally normal^23^. The resulting OddsPath scores were 174 for functionally abnormal variants in our assay, and 0.027 (1/37) for normal variants, corresponding to strong level of evidence towards pathogenicity (4 points) and benignity (−4 points), respectively (^24^; **Table 1**).

**Table 1.** OddsPath-based assay evidence calibration using *RAD51D* clinical variants.

| Pathogenic variants |  |  |  |  | Benign variants |  |  |  |  |
| --- | --- | --- | --- | --- | --- | --- | --- | --- | --- |
| n | Abnormal | Normal | Odds Path | Evidence points | n | Abnormal | Normal | Odds Path | Evidence code |
| 37 | 37 | 0 | 174 | +4 | 174 | 0 | 174 | 1/37 | -4 |

Known pathogenic or benign variants were obtained from ClinVar, requiring two stars (multiple submitters, without discordance), and resolving conflicts when at least two non-VUS interpretations were available (**Methods**). OddsPath scores were determined as described<sup>23</sup>. Variants predicted to be splice disruptive by SpliceAI (score $\geq 0.2$ ) were excluded and treated as ‘not determined’.

We next examined the 789 *RAD51D* missense variants listed in ClinVar as VUS or having conflicting interpretations. We excluded 49 variants predicted to have splicing effects (SpliceAI^25^ scores ≥ 0.2), and the 68 variants that scored as INT and variants that scored as LOF but failed to pass the significance threshold. Of the remaining 672 variants, 87% were functionally normal (NEU); absent any conflicting evidence, the addition of the strong evidence (−4 points) would be enough to reclassify these as likely benign^26^. The other 85 missense variants (12.6%) scored as LOF and could receive 4 points toward pathogenic classification. Nearly two-thirds (n=56) of these were absent from gnomAD, but among those present in that population dataset, the cumulative minor allele frequency (MAF) is 2.03×10^-4^, indicating that, as a lower bound estimate, *RAD51D* missense LOF variants may contribute to hereditary breast and ovarian cancer risk for approximately 1 in 20,000 individuals. As for missense SNVs not yet reported in ClinVar (n=1,264), a similar proportion (∼13%) exhibit abnormal function, and our function scores will provide evidence to classify them as they are newly clinically observed.

### Characterization of *RAD51D* variants identified in breast and/or ovarian cancers for altered DNA repair function through HR and for altered BCDX2 complex formation

To determine how well our variant-function map can predict the ability of *RAD51D* variants to perform HR, we performed a sister chromatid recombination (SCR) assay on 70 clinically-identified breast and/or ovarian cancer *RAD51D* variants, of which 14 were determined to have LOF or intermediate scores in our map (**Supplementary Table 1**). These variants were identified through clinical collaborations, in cancer databases such as Catalog of Somatic Mutations in Cancer (COSMIC) or The Cancer Genome Atlas (TCGA), or in the academic literature (**Supplementary Table 1**). Using the U2OS *RAD51D* KO cells, WT RAD51D or a series of synonymous or truncation alleles were transiently expressed, and the resulting HR activity was used to benchmark and threshold *RAD51D* VUS HR activity (**Supplementary Figure 3; Supplementary Table 2**). To measure HR function, we utilized a GFP-based SCR reporter construct that has been stably integrated into these cells (^19^; **Supplementary Figure 3a**). This assay utilizes a non-functional *GFP*, which is disrupted by the inclusion of a unique *I-SceI* endonuclease cleavage site cloned downstream of a truncated GFP fragment. Upon transfection of a plasmid that expresses the I-SceI enzyme, a DNA double-strand break is induced in the non-functional GFP fragment, which then undergoes DNA repair using the upstream, truncated GFP fragment as a template, thereby restoring GFP expression and subsequent fluorescence. Using this approach, we individually assessed the HR proficiency of 70 breast and/or ovarian cancer-identified *RAD51D* variants relative to known synonymous (benign) or nonsense (pathogenic) variants from the ClinVar database (**Figure 4; Supplementary Figure 3**).

**Figure 4.**
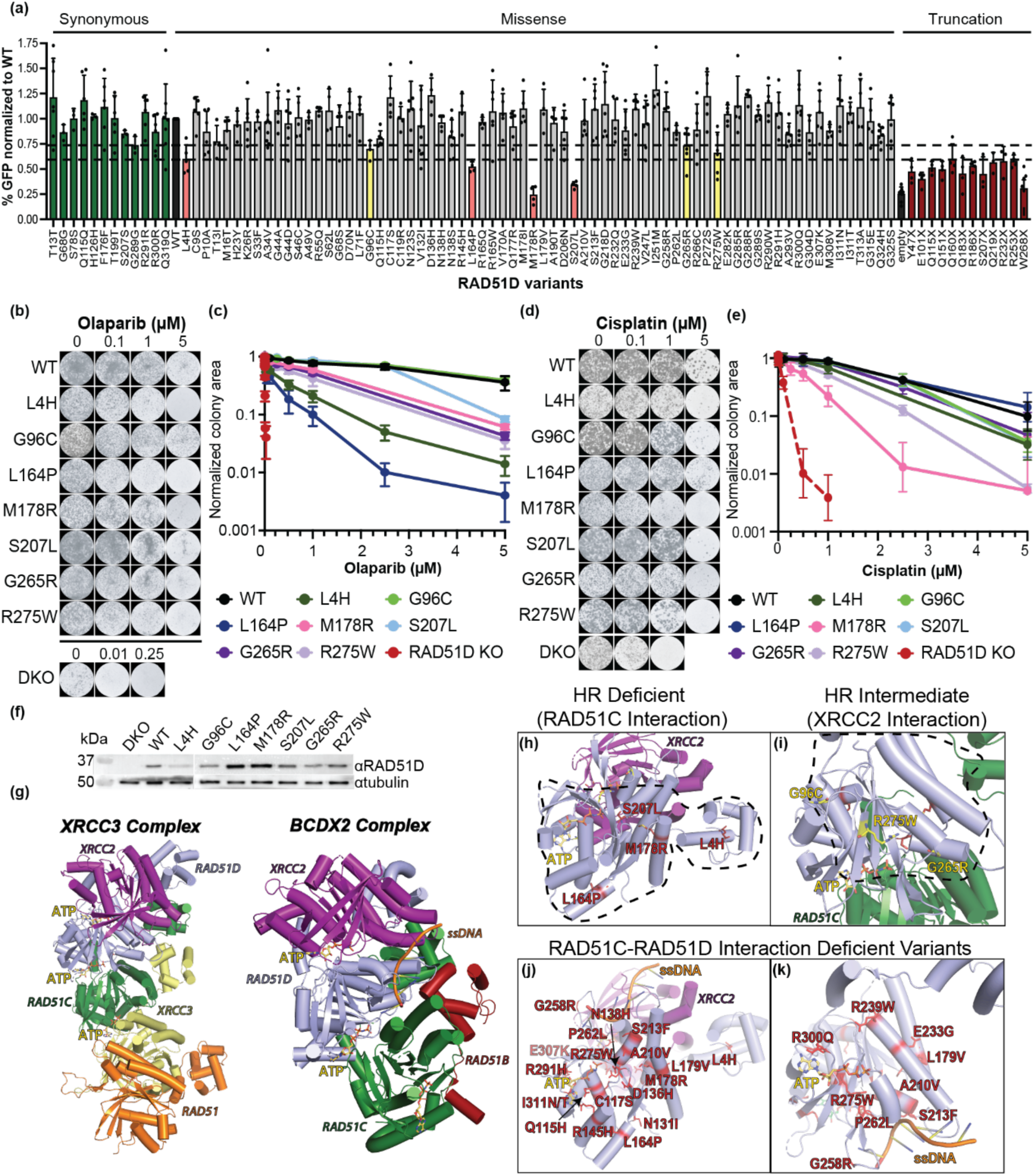
Breast and ovarian cancer-identified variants in the ATPase and DNA-binding regions of *RAD51D* exhibit reduced HR proficiency. **(a)** The HR proficiency of 70 *RAD51D* variants was tested using the sister chromatid recombination assay calibrated against synonymous and truncation variants. Loss of HR function was calculated based on the range of truncation variants (indicated in red) as <0.6 (missense LOF in light). HR proficient variants (gray), were determined based on comparison with the range of synonymous variants (green) if they exhibited HR >0.75. Variants with intermediate HR proficiency, in the range of 0.6–0.75, were color-coded in yellow. See **Supplementary Figure 3b-c** for all synonymous and truncation variants tested, and note that a subset of those variants analyzed are replotted here as representative variants. The experiment was performed 4–9 times and plotted as mean values ± s.d. (**b-e**) Olaparib and cisplatin sensitivity of breast/ovarian cancer identified *RAD51D* variants with reduced HR. Representative images of U2OS cell lines stably expressing WT or the indicated *RAD51D* variant that were treated with increasing concentrations of Olaparib (**b**) or cisplatin (**d**). (**c & e**) Clonogenic survival assays were quantified by percent colony area and normalized to the area of untreated or vehicle control. Means of 4–12 trials are plotted ± s.d., and drug concentrations with colony area <0.001 are omitted. (**f**) *RAD51D* variants with reduced HR are expressed. Western blot analysis of U2OS cell lines stably expressing WT or the indicated *RAD51D* variants. RAD51D protein expression was assessed using an anti-RAD51D antibody, and equal protein loading was assessed using an anti-Tubulin antibody. Note that L4H exhibits reduced protein expression. Experiment performed in triplicate. (**g**) Structures of the RAD51 paralog pentamer, XRCC3 complex, (PDB: 9SVX) and the BCDX2 complex (PDB: 8GBJ). Cartoon representations of XRCC2 (purple), RAD51D (light blue), RAD51C (green), XRCC3 (yellow), RAD51 (orange), and RAD51B (red) are shown consistently throughout. Bound ATP molecules are represented as yellow-orange sticks. ssDNA is shown in orange. (**h-i**) Variants causing complete (red, **h**) or intermediate (yellow, **i**) deficiency in homologous recombination are highlighted on RAD51D. The interacting surfaces of RAD51C and XRCC2 are outlined with black dashed lines. The contact interfaces are identical between the XRCC3 and BCDX2 complex. (**j-k**) Variants that disrupt RAD51C-RAD51D interactions are shown as red sticks, mapped onto the BCDX2 complex structure.

We observed a strong correlation with MAVE results (Pearson’s *r*: 0.75, P=1.27×10^−19^; **Figure 5d**) and identified four *RAD51D* HR-deficient variants (L4H, L164P, M178R, S207L) and three alleles with attenuated HR (G96C, G265R, R275W) (**Figure 4a**, **Supplementary Table 1**). We verified expression of the RAD51D missense variants, and all variants are expressed, although to varying degrees (**Supplementary Figure 4a**). Since these HR-disrupted variants all had LOF scores in the MAVE olaparib selection assay, using a clonogenic survival assay, we sought to validate their sensitivity to olaparib and test whether they exhibited sensitivity to cisplatin, which is a platinum-based compound used as a front-line treatment for ovarian cancer^27^ (**Figure 4b-e**). To do this, we created stable cell lines in the U2OS *RAD51D* KO cell lines solely expressing the variant of interest and validated both expression and loss of HR function compared to our transiently transfected cell lines (**Fig 4f; Supplementary Figure 4b**). Using these stably complemented cell lines, we found that cells expressing the HR deficient variants exhibited variable sensitivity to olaparib and/or cisplatin, but none were as sensitive as the *RAD51D* knockout cell line (**Figure 4b-e**).

**Figure 5.**
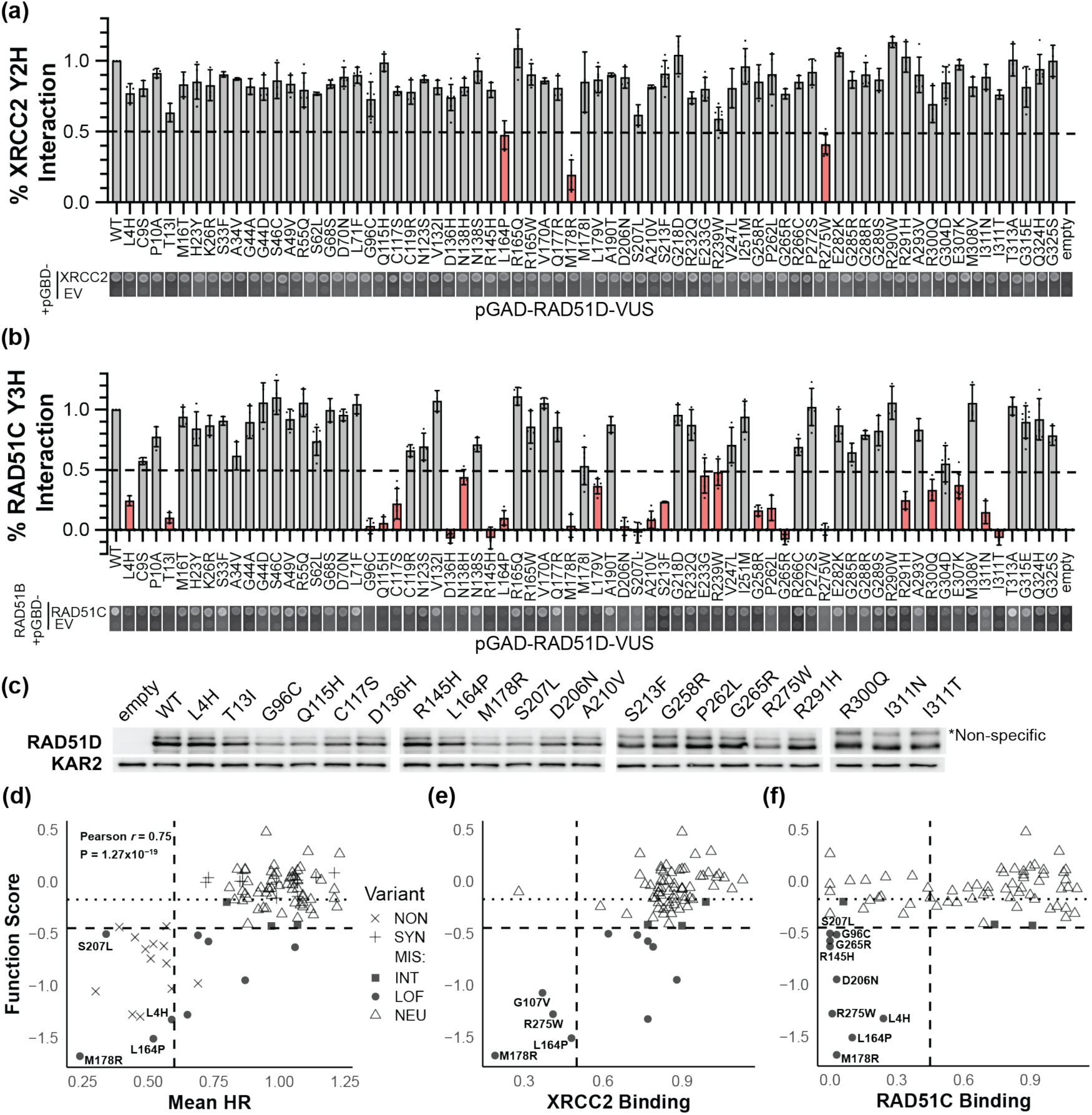
A subset of RAD51D breast and ovarian cancer identified variants alter interaction with XRCC2 or RAD51C by Y2H or Y3H, respectively. (**a-b**) Interaction proficiency of RAD51D VUS was measured via (**a**) yeast 2-hybrid (Y2H) of pGAD-RAD51D or its variant, expressed in a GAL4 DNA activating domain expressing plasmid with pGBD-XRCC2 expressed in the GAL4 DNA binding domain expressing plasmid and (**b**) pGAD-RAD51D or its variant, expressed in the pGAD (GAL4 DNA activating domain) plasmid with pGBD-RAD51C (GAL4 DNA binding domain) plasmid with pADH1-RAD51B via yeast 3-hybrid (Y3H). RAD51B serves to stabilize RAD51C protein levels. Empty vectors are used as a negative control. Quantification of yeast growth from three experiments is plotted as mean ± s.d relative to the WT control. Representative images for variants and controls are shown. Variants with <50% interaction are classified as deficient and plotted in red. (**c**) RAD51D variants with reduced interaction with XRCC2 or RAD51C are expressed. Western blot analysis of protein extract from yeast cells in (a) expressing WT RAD51D or the indicated RAD51D variants was assessed using an anti-RAD51D antibody, and equal protein loading was assessed by measuring the amount of Kar2 using an anti-Kar2 antibody. Note that a subset of variants has reduced protein expression relative to WT RAD51D. Experiment performed in triplicate. See **Supplementary Figure 5** for Western blots of all variants. (**d-f**) MAVE function scores vs individual variant assay results of **(d)** HR activity, **(e)** XRCC2 binding, and **(f)** RAD51C binding. Missense variant point shape denotes functional status (LOF/INT/NEU) based upon MAVE result.

Upon mapping the identified HR deficient variants on the solved cryo-EM structures of RAD51D within the BCDX2 and X3CDX2 complexes, we observed no clear localization of the critical residues: two of the HR deficient residues are located in the ATP-binding core (L164 and S207), two are found in the DNA-binding loop (G265, R275), while M178 is located in the RAD51D-RAD51C binding interface, and L4 in the N-terminus (**Figure 4h-i**). Together, these results suggest that variants that disrupt the ATPase core domain or DNA binding interfaces may be critical for HR function.

RAD51D is a member of the BCDX2 and X3CDX2 complexes, where it directly interacts with XRCC2 as part of the subcomplex DX2 and with RAD51C, which is part of the BC or X3C subcomplex^11,28^. Our previous work on RAD51C demonstrated that variants that disrupt BCDX2 complex formation are indicative of loss-of-function^15,16^. To determine whether any of the 70 breast/ovarian cancer identified variants alter BCDX2 complex formation, we examined the RAD51D variants interaction with XRCC2 by yeast-2-hybrid analysis (Y2H). Using this approach, we found that three of the variants had reduced interaction with XRCC2 (**Figure 5a**; L164P, M178R, R275W), and all exhibited LOF (**Figure 5e**). We next examined whether the RAD51D variants would alter their interaction with RAD51C using a yeast-3-hybrid system (Y3H), where RAD51B is used as a third vector to help stabilize RAD51C. Importantly, 26 of the 70 RAD51D variants exhibited defects in interaction with RAD51C and had an overall intermediate MAVE score, with 10 functionally defective variants (mean=-0.940), including the three variants that also exhibited loss of XRCC2 interaction (**Figure 5e-f**). Expression of the variants from the yeast cells was confirmed by Western blot (**Figure 5c; Supplementary Figure 5**). In some cases, the defect in protein interaction could be due to reduced expression (**Supplementary Figure 5; S46C**). Together, these findings suggest that the interaction between RAD51D and its binding partners, XRCC2 and RAD51C, is critical for function.

To determine if loss of HR observed in the RAD51D cancer-identified variants results in alteration in complex formation or RAD51D biochemical activity, we purified recombinant DX2 subcomplex with or without one of the seven different RAD51D variants with a FLAG-tagged XRCC2. We then incubated the BC subcomplex with the DX2 subcomplex and performed pulldown assays using DX2-FLAG. We found that four of the RAD51D variant-containing complexes exhibited defects in BCDX2 complex formation (**Figure 6a**; G96C, L164P, M178R, S207L, as indicated by the comparison of enrichment in the eluate to the flow-through). To further investigate the mechanisms driving LOF in these variants, we performed functional analysis using these complexes using BCDX2 complexes reconstituted by mixing WT BC and DX2 subcomplexes with later containing either WT or mutant RAD51D.

**Figure 6:**
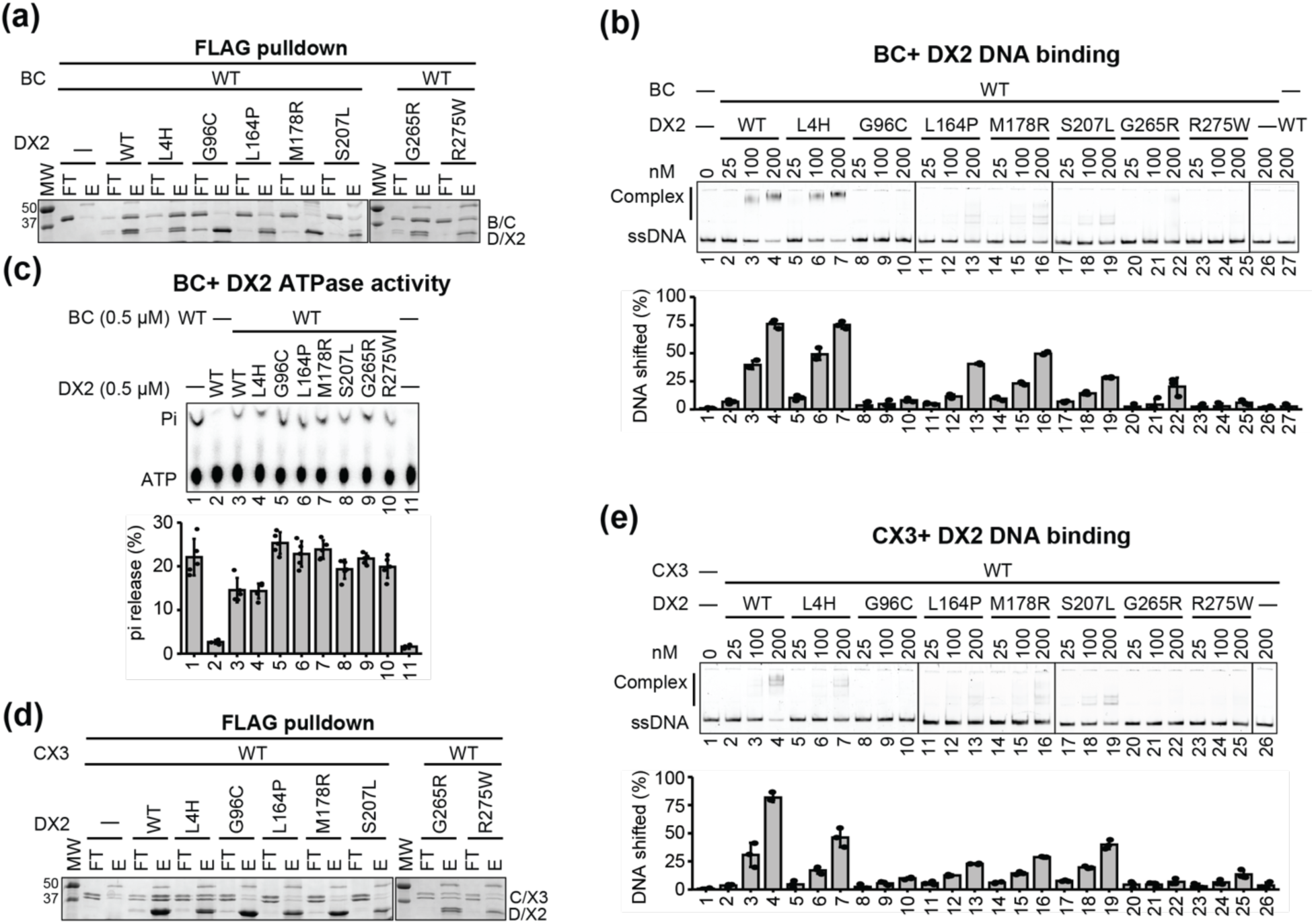
RAD51D HR deficient variants differentially impact ssDNA binding, BCDX2 ATPase activity, and/or RAD51-mediated strand exchange. (**a**) Pull-down analysis of WT BC (RAD51B-His/RAD51C) and DX2 (RAD51D/XRCC2-FLAG) sub-complexes with indicated RAD51D variants using anti-FLAG resin. (**b**) ssDNA binding of BCDX2 paralog complexes reconstituted by mixing WT BC and DX2 sub-complexes with indicated RAD51D variants. (**c**) Pull-down analysis of WT CX3 (RAD51C-His/XRCC3-STREP) and DX2 (RAD51D/XRCC2-FLAG) sub-complexes with indicated RAD51D variants using anti-FLAG resin. (**d**) ATPase analysis of 0.5 μM WT BC and DX2 sub-complexes with indicated RAD51D variants in the presence of ssDNA after 60 min incubation. Pi indicates released inorganic phosphate after hydrolysis. (**e**) ssDNA binding of X3CDX2 paralog complexes reconstituted by mixing WT CX3 and DX2 sub-complexes with indicated RAD51D variants. For (**b, d-e**), results from three independent experiments were plotted as mean values ± s.d.

### Six of the seven RAD51D variants with reduced HR exhibit altered DNA binding and BCDX2 ATPase activity

The association of BC and DX2 sub-complexes into the BCDX2 complex synergistically enhances ssDNA binding at DNA break sites to facilitate RAD51 filament formation in an ATPase-dependent manner^12,16^. To assess whether RAD51D variant can affect ssDNA binding of BCDX2 complex, we conducted electrophoretic mobility shift assays (EMSA) in the presence of Mg²⁺/ATP using an 80-nucleotide single-stranded DNA substrate at increasing concentrations BCDX2 complexes (**Figure 6b**). BCDX2 complex containing RAD51D-L4H is DNA binding proficient, whereas six of the seven RAD51D variants displayed attenuated DNA binding compared to WT BCDX2 (**Figure 6b**). We note that defective DNA binding of BC with DX2 sub-complexes containing RAD51D variants G96C, L164P, M178R, S207L was primarily attributable to defective BCDX2 complex assembly (**Figure 6a-b**). In contrast, RAD51D variants G265R and R275W formed intact BCDX2 complexes but displayed severe DNA-binding deficiencies, consistent with their mutations residing within the second DNA-binding loop of RAD51D (**Figure 6b and Figure 4j-k**).

Our previous work demonstrated that the BC sub-complex exhibits robust ssDNA-dependent ATPase activity, whereas the DX2 sub-complex does not. Interestingly, addition of DX2 to BC reduced ATP hydrolysis, suggesting that DX2 modulates BC ATPase activity at DNA break sites^12^. Next, we examined whether the regulation of BC ATPase activity by DX2 would be altered in complexes containing the various RAD51D variants. Similar to wild type, DX2 with RAD51D-L4H dampened BC ATP hydrolysis (**Figure 6c**). In contrast, six variants defective in ssDNA binding failed to influence BC ATPase activity, as ATP hydrolysis remained comparable to BC alone (**Figure 6c**). These findings indicate that the synergistic stimulation of ssDNA binding by BC-DX2 association is essential for ATPase regulation within the BCDX2 complex, and that RAD51D variants impaired in complex formation and/or ssDNA binding cannot modulate BC ATPase activity.

### Identification of RAD51D-L4H as a separation-of-function mutant proficient for BCDX2 activity with reduced X3CDX2 function

It was recently discovered that in addition to RAD51D’s central function in the BCDX2 complex, the DX2 subunit is a member of another RAD51 paralog-containing complex with RAD51C and XRCC3, X3CDX2^11^. Notably, interfaces of RAD51C and RAD51D interaction are similar in both BCDX2 and X3CDX2 complexes^11,12,29^. Therefore, we asked whether RAD51D variants with altered HR would also exhibit defects in X3CDX2 formation. To address this question, we examined DX2 subcomplexes containing WT RAD51D, or the variant of interest, for interaction with CX3 (**Figure 6d**). Similar to BC–DX2 interactions, CX3 associates with DX2 when RAD51D contains L4H or G265R, but not with variants G96C, L164P, M178R, or S207L (**Figure 6d**). Additionally, unlike BC–DX2, CX3 fails to interact with DX2 carrying RAD51D-R275W (**Figure 6d**). We then assayed whether ssDNA binding of the X3CDX2 complex would be impacted, and indeed, all the tested variants exhibit reduced ssDNA binding or form some lower molecular weight complexes (**Figure 6e**). It is important to note that while RAD51D-L4H exhibits no discernible biochemical defect within the BCDX2 complex and still forms the X3CDX2 complex, it shows attenuated DNA binding within the latter (**Figure 6e**). These results suggest that RAD51D-L4H is a separation-of-function allele that may uncouple the activities of the BCDX2 and X3CDX2 complexes.

## Discussion

To identify critical regions for RAD51D function, we undertook a high-throughput screen of RAD51D’s response to olaparib, examining 6,888 synonymous, nonsense, and missense changes across the full-length *RAD51D* cDNA. We identified critical regions near ATPase-like domains, as well as the binding interfaces between RAD51C-RAD51D and RAD51D-XRCC2, and key structural elements critical for olaparib response. Functional screening of 70 clinically identified variants for HR and/or BCDX2 complex interactions showed high correlation with our MAVE results, where HR-deficient variants and variants that disrupt RAD51D interaction with XRCC2 were also significantly depleted. As RAD51D variants are routinely uncovered in high-risk patients for breast and ovarian cancer, these findings validate and enforce the utility of using high-throughput variant analysis and classification.

While RAD51D’s role as a tumor suppressor is well established, surprisingly little is known about its biochemical function during HR. Our groups recently found that RAD51D, when partnered with XRCC2, attenuates BC ATPase activity^12^. The significance of this attenuation has been largely overlooked. Here, we provide evidence that RAD51D DNA-binding and ATPase-like domains are critical for this regulation (summarized in **Figure 7a-b**). These results are unexpected, given that the BCDX2 complex is an ATPase, and one might have expected that DX2 would stimulate ATP hydrolysis of BC, rather than decrease it. Clues to this regulatory mechanism come from RAD51 itself. RAD51 is an ATPase that forms nucleoprotein filaments on the DSB ends, and its affinity for DNA is strongly enhanced upon ATP binding, whereas ATP hydrolysis results in RAD51 removal from the ssDNA^30^. Several key regulatory factors rely on this mechanism to regulate RAD51 loading at sites of DNA damage, such as Srs2 in yeast^31–33^ and REQCL5 and FIGNL1 in humans^34–36^. Though BC subcomplex displays weak ssDNA binding, like RAD51, its ATPase activity is stimulated upon ssDNA. In contrast, despite having ATPase core domains and motifs, DX2 is not an ATPase; rather, its interaction with BC attenuates this ATPase activity and is required for stable ssDNA binding of BCDX2 complex^12^. Given the high degree of conservation between RAD51 and its paralogs, we hypothesize that slowing the hydrolysis of ATP would increase BCDX2 DNA binding dwell times and therefore facilitate RAD51 filament elongation (**Figure 7c**). In contrast, loss of DX2 or its DNA-binding capabilities results in increased ATP hydrolysis of BC, thus resulting in reduced RAD51-mediated HR. Indeed, our most severe HR-deficient variants impact this ATPase regulation, and all decrease DNA binding to various degrees.

**Figure 7:**
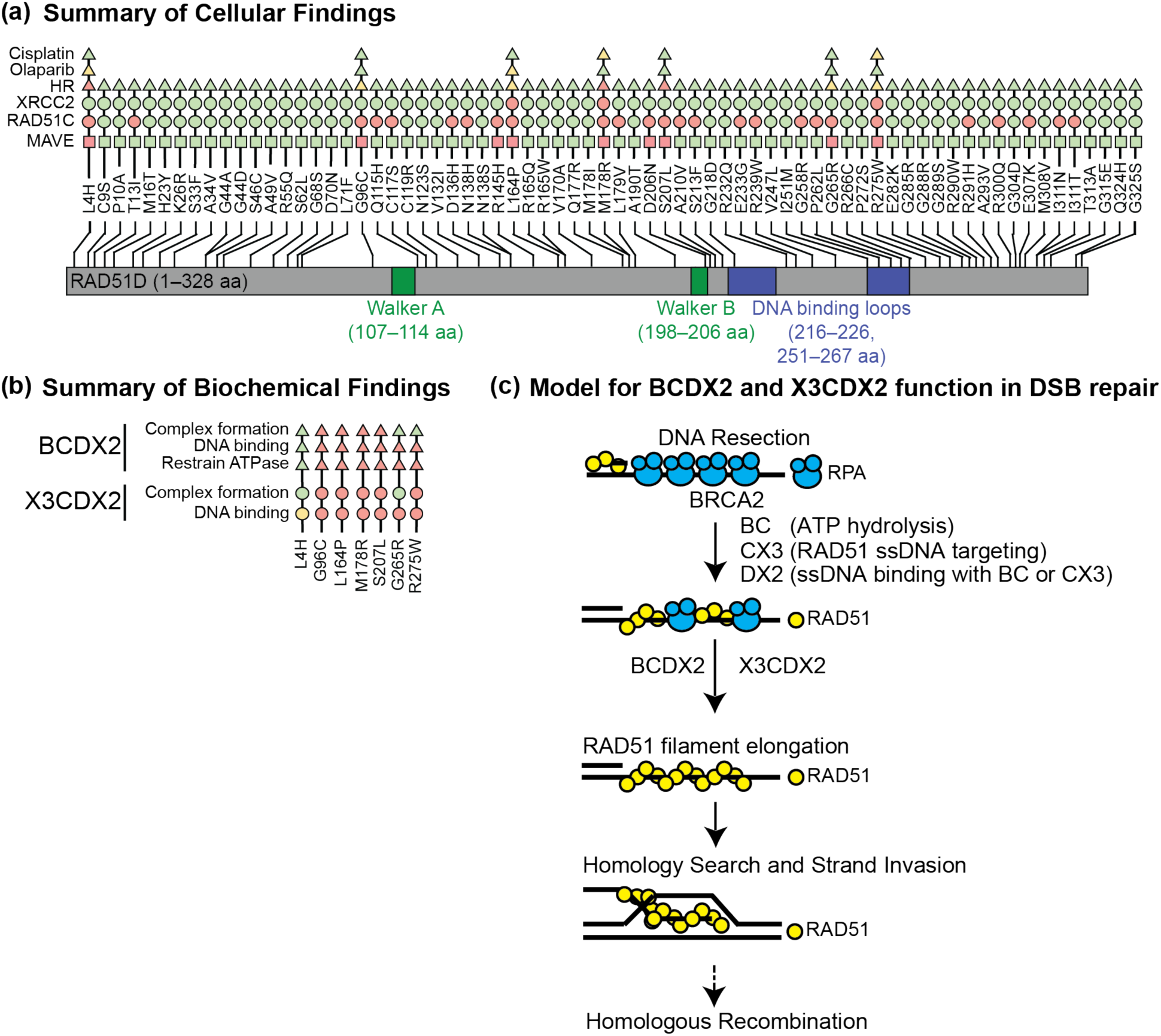
Summary of findings and model for RAD51D function as part of the BCDX2 and XRCC3 (X3CDX2) complexes. (**a**) Summary of cellular findings. Schematic of RAD51D (1-328 aa) is shown with the Walker A and B motifs (green), and DNA binding loops (blue) indicated. The variants analyzed are shown above with the results from the cellular studies (MAVE variant-function map; Figure 2), RAD51D Y2H interaction with XRCC2 and its Y3H interaction with RAD51C (Figure 5), SCR recombination (HR), as well as olaparib and cisplatin clonogenic survival assays (Figure 4) are summarized based on functional score. Loss-of-function is shown in red, an intermediate function is shown in yellow, and wild-type function is indicated in green. (**b**) Summary of biochemical findings. Variants included in biochemical analysis for BCDX2 or XRCC3 (X3CDX2) complex formation, DNA binding, and ATPase restrain from Figure 6 are shown. (**c**) Model for BCDX2 and X3CDX2 function in DSB repair. Upon DSB formation, the 3’ ssDNA end is resected and coated by RPA. The BCDX2 and X3CDX2 complexes facilitate the displacement of RPA and the loading of RAD51 onto ssDNA. This is achieved by the regulation of BC ATPase activity by DX2, which facilitates its binding to ssDNA. CX3 in complex with DX2 then targets RAD51 to ssDNA. The combined functions of the BCDX2 and X3CDX2 complexes are necessary to facilitate RAD51 filament assembly and subsequent RAD51-mediated homology search and strand exchange activities.

MAVE analysis is useful not only for ascertaining the novel variants’ impact, but also its global view of functional constraint has enabled key mechanistic insights into the structure and function of RAD51D. When we mapped the depleted variants of RAD51D onto the recently solved cryo-EM structure of BCDX2 complexed with DNA, we found five beta sheets that are aligned within the central structure of RAD51D, which, when mutated, result in loss of function (**Figure 3**). Our high-resolution functional mapping of this structure showed high intolerance to mutation, pairing well with structural data, and together suggests that this core RAD51D beta sheet is required to promote this modulatory effect—possibly by propagating conformational changes based on the RAD51D-DNA interface (**Figure 3f**). Furthermore, our Y3H experiments indicate that the RAD51C-RAD51D interaction is much more fragile than the RAD51D-XRCC2 interaction, with many protein-coding changes fully disrupting the BC-DX2 complex. Further biochemical and structural studies into critical structural regions of RAD51D and mechanisms driving its modulatory functions are warranted.

High-throughput functional assays can assist clinical variant interpretation, provided their measurements are carefully evaluated and calibrated^6,37^. Supporting our map’s suitability for clinical use, it perfectly separated synonymous variants from nonsense variants (excluding truncations after codon 305). Likewise, classifications based upon our function scores and cutoffs were completely concordant with the standing classifications across the 31 missense variants with consensus interpretations in ClinVar. Calibrated to these clinical variant sets, our map meets the ‘strong’ level of evidence in both directions of effect (i.e., for or against pathogenicity) under accepted clinical variant interpretation guidelines^23^. Thus, our map will assist the resolution of 789 existing missense VUS in *RAD51D* and inform the interpretation of the other 1,264 such variants as they are detected clinically in the future.

Our MAVE analyses are also corroborated by orthogonal functional assays, including HR and protein interaction assays for 70 individually tested variants. We further compared our function scores to a saturation genome editing (SGE) map of *RAD51D* (^38^; *Casedei et al., in preparation*). These two maps are highly significantly correlated (Pearson’s *r*=0.58, P<10^-226^), and yielded consistent classifications for 92.9% of the variants scored by both (**Supplementary Figure 6**). Among these, for the 270 of these listed in ClinVar with two or more non-VUS interpretations, the two maps agreed with each other and with the ClinVar consensus interpretation in 263 cases (97.4%). This level of concordance is remarkable for maps produced by separate groups using different cell lines (U2OS vs HAP1) and distinct functional selections (olaparib selection vs cell fitness).

Nevertheless, a potential limitation of our cDNA-based MAVE is that it cannot capture variant effects upon mRNA splicing. We used SpliceAI to flag 120 variants with potential to disrupt mRNA splicing despite having normal protein activity in the context of an unspliced cDNA; we removed these from our comparison to clinical variants and did not apply functional evidence to them for clinical variant classification. This filter captures nearly half of the variants (41/92) that appeared neutral in our assay but were abnormal in SGE, which unlike our approach, can detect loss of function due to splicing disruption. Conversely, our cDNA-based approach detected a broad range of intermediate effects which may on their own be too subtle to perturb cell fitness in SGE and related assays. As it remains unclear whether these hypomorphic variants are pathogenic, we did not use them to apply clinical evidence. Even so, the functional constraint they reveal enabled our new insights into RAD51D function. Likewise, by interrogating the full missense space, our map highlights the constraint patterns such as the complete constraint of glycine 265, and the strong preference for compact hydrophobic residues at leucines 4, 12, and 20.

One important finding from our work is that most clinically identified variants have little to no impact on HR. While it is possible that the majority of these variants are passenger mutations or benign, it suggests that other additional functions of RAD51D may also be critical for tumor suppression. For example, RAD51D has an HR-independent function in replication fork progression; however, the significance of this function in cancer is unknown. The modulatory role of RAD51D may serve critical additional functions to promote RAD51 filament formation, depending on the DNA substrate and cell cycle contexts, thereby facilitating high-fidelity DNA repair. Furthermore, our screening methodologies may not detect variants with replication defects. Future studies examining this important aspect of DNA repair will shed light on the impact of BCDX2 and X3CDX2 replicative functions in promoting genomic integrity and cancer prevention.

## MATERIALS AND METHODS

### RAD51D Directed Mutagenesis Screen

#### Tissue Culture, Cell lines, and Reagents

Human osteosarcoma U2OS SCR (sister chromatid recombination) #18 wild-type (gifted from Mauro Modesti;^19^) and *RAD51D* CRISPR knockout (KO; clone #4, purchased from DSMZ, no. ACC835) were cultured in Corning DMEM supplemented with 10% fetal bovine serum (FBS) and penicillin-streptomycin antibiotic (50 U/mL). Cells were incubated at 37°C in a 5% CO_2_ atmosphere. Cells were transfected using Lipofectamine 2000 (Invitrogen) diluted with OptiMEM serum-free media following the manufacturer’s instructions. Cisplatin and olaparib (Selleck Chem; AZD2281).

#### Library synthesis and transduction

Saturation mutagenesis libraries were prepared as described^39,40^; briefly, a wild-type *RAD51D* cDNA plasmid (GenBank: NM_002878.4) was synthesized (Twist Bioscience). The cDNA was divided into 6 tiles (**Supplementary Table 3**), and for each, a mutant oligonucleotide library was designed in which each codon was sequentially replaced by an ‘NNN’ randomer. Mutant oligonucleotides were synthesized together as a single oPool oligo pool (IDT), then PCR-amplified using tile-specific primers to add sequences for cloning into the starting cDNA plasmid. Mutant cDNA libraries were cloned into a ‘Tet-On’ inducible lentiviral expression construct (pCW57.1; Addgene #80922), transformed into Endura electrocompetent cells (Lucigen). Plasmids were prepared with ZymoPure II kits, and pooled lentivirus libraries were prepared by the University of Michigan Vector Core. U2OS *RAD51D* knockout cells were transduced with WT *RAD51D* lentivirus or pooled mutant lentivirus, at low multiplicity of infection (MOI<0.1) such that each cell carries zero or one stably integrated copies of *RAD51D* cDNA. Per replicate, three million cells were infected with 200 µL lentiviral supernatant in the presence of 8 μg/mL polybrene. At 48 hr post-infection, transduced cells were selected with 50 µg/mL hygromycin, which was maintained in all subsequent steps.

#### Olaparib functional selection assay

Mutant library-expressing cells were cultured in experimental triplicates in T175 flasks in DMEM media (supplemented with 10% fetal bovine serum and 1% penicillin-streptomycin) with 1 mg/mL doxycycline added to induce *RAD51D* expression. From each baseline passage (‘‘P0’’) culture, one million cells were collected for gDNA extraction and three million cells were seeded for olaparib selection, with media containing 50 µg/ml hygromycin, 1 mg/mL doxycycline, 1 µg/mL blasticidin (to enforce *RAD51D* expression), and 250 nM olaparib (as used in ^41^). Cells were grown to 90% confluence for two rounds. At each round, three million cells were harvested for gDNA extraction and then seeded with three million cells to avoid population bottlenecks.

#### Mutant library sequencing

Genomic DNA from P0 and P2 cell populations was isolated using CleanNA Blood and Tissue magnetic bead kit (Bulldog Bio). To amplify integrated *RAD51D* mutant libraries, “outer PCR” was performed on a 500 ng gDNA template with primers flanking the *RAD51D* cDNA using the PrimeSTAR GXL kit (Takara), with 16 replicate PCR reactions for each gDNA sample to maintain library complexity. Gel electrophoresis and shallow shotgun sequencing of the full-cDNA PCR products were performed to confirm the absence of large rearrangements, insertions, or deletions across cell passages.

To quantify mutation frequency in sorted cells, mutant libraries were subjected to deep amplicon tile sequencing^42^ as described^39^. Briefly, 1 µL of outer PCR library product was amplified in a limited (≤10) cycle PCR, using primers directed at constant bases flanking the targeted tile within the *RAD51D* cDNA. Primer 5’ tails contained constant sequencing adaptor stub sequences. Each resulting product was then diluted and further amplified to add sequencing adaptors and sample-specific unique dual 10 bp index sequences. Amplicon size was verified by electrophoresis on 6% TBE native PAGE gel stained with SYBR Gold. Indexed libraries were pooled, purified with SPRI beads, and sequenced with paired-end 150 bp reads on an Aviti24 sequencer (Element Biosciences) at the University of Michigan Advanced Genomics Core. Libraries were sequenced to a median depth of 12 million read pairs per replicate.

#### Function score calculation

Paired-end reads were overlapped, and an error-corrected consensus was taken by PEAR^43^, then aligned to a *RAD51D* cDNA reference with bwa mem^44^. Variant counts were tabulated by a pipeline implemented in Snakemake (https://github.com/kitzmanlab/tileseq/). Counts were summed across synonymous codons and the resulting amino acid-level counts for each replicate were processed by Rosace^45^, yielding a single ‘fitness’ score for each variant and a local false sign rate (lfsr) score, a measure of confidence that the sign of the fitness score is correct^46^ (**Supplementary Tables 4,5**). To normalize between tiles, each variant score *s_i_* was scaled and shifted to center synonymous scores at 0, and nonsense scores at −1: *r_i_’*=(*r_i_* -med_syn)/(med_syn-med_non), where med_syn and med_non are per-tile medians of *r_i_* for synonymous and nonsense variants, respectively. Variants with low initial abundance (≤100 counts per million at timepoint P0; n=154, 2.2%) were excluded from further analysis. For DE intolerance analysis, function scores were scaled with minimum score = 0, then for each residue, the mean score of DE variants was subtracted from the mean score of all non-DE variants.

#### Variant classification

We determined functional cutoffs by fitting normal distributions to the scores of nonsense variants (residue ≤304) and synonymous variants, and taking the 97.5^th^ percentile and 2.5^th^ percentile of the respective distributions. Variants scoring below the lower cutoff (−0.446) and having an lfsr ≤0.01 were classified as loss-of-function (LOF). Variants scoring above the upper cutoff (score ≥-0.170) were considered as neutral (NEU). Variants with function scores between the two cutoffs and with lfsr ≤0.01 were classified as intermediate (INT), and these were not used for clinical variant analyses. Variants in the intermediate scoring range but with lfsr >0.01 were considered as neutral. Variants with LOF scoring but lfsr >0.01 were excluded from analysis (n=38, 0.57 %). ClinVar interpretations were downloaded on 9/28/2025. Variants with ‘soft conflicts’ were included only if there were two or more non-VUS interpretations, all in the same direction (e.g., all B/LB or all P/LP). Splicing effects were predicted by SpliceAI^25^, and variants with deltaMax score ≥0.2 were considered be potentially splicing-disruptive.

#### RAD51D missense VUS selection

RAD51D missense variants were identified in databases (COSMIC, TCGA, ClinVar), as well as through our clinical collaborators and the literature (**See Supplementary Table 1**). Of the missense RAD51D variants in our list, 70 variants were previously reported in ClinVar as somatic or germline variants identified in breast and ovarian cancers and either CON/P or VUS (with the exception of p.Ser207Leu (S207L) classified as P/LP and p.Arg165Gln (R165Q), p.Arg232Gln (R232Q), p.Glu233Gly (E233G) classified as B or B/LB).

#### Molecular cloning

All plasmid constructs were prepared through either site-directed mutagenesis of the corresponding *RAD51D*-WT cDNA template or synthesized by Gene Universal (Gene synthesis and mutagenesis services, Newark, DE). For yeast assays, *RAD51D*-WT was cloned into the pGAD-C1 backbone at the 5’-*EcoRI* and 3’-*SalI* restriction cut sites. Breast and ovarian cancer-identified VUS were created by site-directed mutagenesis using primers (synthesized by Integrated DNA Technologies) that contained the corresponding nucleotide mutation. For yeast assays, *RAD51C* and *XRCC2* were cloned into the pGBD-C1 backbone at the 5’-*EcoRI* and 3’-*SalI* restriction cut sites. For cell-based assays, *RAD51D*-WT and corresponding mutated VUS were subcloned into the 5’-*HindIII* and 3’-*EcoRI* restriction sites of the cFlag pcDNA3 backbone. cFlag pcDNA was a gift from Stephen Smale (Addgene plasmid #20011). All mutations were validated through Sanger sequencing (Azenta).

#### Yeast-2-hybrid and yeast-3-hybrid analysis

Yeast-2-hybrid (Y2H) and yeast-3-hybrid (Y3H) assays were performed as previously described^20,47^, with the exception that *RAD51D* WT and corresponding VUS were subcloned into the pGAD-C1 backbone (as described above) and expressed as C-terminal fusions to the GAL4 activating domain. Wild-type *XRCC2* and *RAD51C* were subcloned into the pGBD-C1 backbone (as described above) and expressed as a C-terminal fusion to the GAL4 DNA binding domain for Y2H and Y3H, respectively. Following co-transformation in the PJ694-alpha yeast strain, 3-4 single colonies were cultured in liquid SC-Leu^−^Trp^−^ (Y2H) or SC-Leu^−^Trp^−^Ura^−^ (Y3H) media overnight. Plates were incubated for 72 hr at 30°C before imaging. Colony growth was quantified in ImageJ and normalized to the respective wild-type (positive) and empty vector (negative) controls. Images were adjusted for uniform brightness and contrast using Adobe Photoshop.

#### Homology-directed repair reporter assay

Plasmid constructs of each *RAD51D* VUS were subcloned into the cFlag pcDNA3 backbone (Addgene #20011) as described above. pCBA-SceI was provided as a gift from Maria Jasin (Addgene plasmid #26477). U2OS-SCR18-WT and U2OS-SCR18-RAD51D-KO cells were seeded in a single well of a 6-well plate at densities of two and four million cells per well, respectively (due to slower growth rate of *RAD51D* knockout cells). Cells were incubated for 12-24 hr before liposomal-mediated transfection with pCBA-SceI (2 µg) alone or co-transfected with empty vector (cFlag pcDNA3, negative control) or vector expressing individual *RAD51D* VUS (2 µg, 1:1 ratio) for 4 hr. Transfection efficiency was assessed by transfecting a single well with pEGFP-N1. Following a 48 hr incubation, cells were collected by trypsinization and fixed in an aqueous 2% paraformaldehyde solution. Fixed cells were re-suspended in phosphate-buffered saline (PBS) and analyzed on the BD Accuri C6 Plus flow cytometer. For each sample, 30,000 events were recorded, and results were normalized to the complemented *RAD51D* WT control. Each variant was analyzed in at least three biological replicates.

#### Western blot

For analysis of protein levels in yeast, 3-4 single colonies of PJ69-4α co-transformed with pGAD-RAD51D-WT or VUS and pGBD-XRCC2 were cultured overnight. The culture density was measured by spectrophotometry (0.75 OD_600_) and yeast were collected through centrifugation and proteins isolated through TCA precipitation. For the analysis of protein levels in mammalian cells, cell lysates were prepared by lysing at least 4 million cells in RIPA buffer. Protein concentration in cell lysates was measured against a BSA standard curve (0-2.0 mg/mL) using the BCA protein quantification kit. Yeast (25 µL) and cell lysates (10 µg) were resolved on 10% SDS-PAGE gels with 4% stacking gels. Proteins were transferred to PVDF membranes and blocked with either 5% [w/v] milk or 5% [w/v] BSA diluted in Tris-buffered saline supplemented with 1% Tween-20 (TBST, blocking buffer is antibody specific). Antibodies against RAD51D (Abcam, 202063, 1:1000), KAR2 (Santa Cruz, sc-33630, 1:1000), and α-tubulin (Cell Signaling Technologies, 2144, 1:1000) were diluted in their respective blocking buffers and incubated with the membrane at 4°C overnight. The next day, membranes were rinsed with TBST and then incubated with secondary antibody (Jackson anti-rabbit #111-165-003, 1:10,000) diluted in TBST for 1 hr. Yeast blots were developed on film. The chemiluminescence reagent was imaged on a Bio-Rad Gel Doc with exposure times ranging from 5 to 300 seconds.

#### Clonogenic survival assays

U2OS-SCR18-WT and U2OS-SCR18-RAD51D-KO cells stably complemented with *RAD51D* WT or *RAD51D* VUS were seeded in a 6-well plate in DMEM supplemented with FBS (10%) and penicillin-streptomycin antibiotic (50 U/mL) at cell densities ranging from 1,000 to 5,000 cells per well (for cisplatin treatment) and 10,000 to 20,000 cells per well (for olaparib treatment). For cisplatin treatment, cells were incubated in media supplemented with cisplatin (at least nine unique treatment concentrations ranging from 0.05 nM to 25 µM) for 48 hr, at which point cisplatin-supplemented media was removed and replaced with fresh, non-treatment-supplemented media. Cells were incubated for 14 days post-cisplatin treatment, and the media was refreshed 7 days post-treatment. For PARPi treatment, cells were incubated in either DMSO- (0.1%) or olaparib- (at least nine unique treatment concentrations ranging from 0.01 nM to 25 µM) supplemented media. Cells were incubated for a total of 15-21 days following initial treatment, and DMSO or PARPi-supplemented media were refreshed every 3-4 days. At the end of incubation, the media was aspirated, and the cells were rinsed with PBS. They were then fixed in 100% methanol and stained with crystal violet. Cells were imaged on a BioRad ChemiDoc Imager, and colony area was quantified using the colony area plug-in tool in ImageJ. All variants and controls were analyzed in at least three biological replicates.

#### Protein expression and purification

Purifications of DX2 (RAD51D and XRCC2-FLAG) and BC (RAD51B-His and RAD51C) sub-complexes were carried as previously described^12^. For purification of the CX3 (RAD51C-His and XRCC3-Strep), Hi5 insect cells were infected for 48 hrs with baculoviruses expressing RAD51C-His and XRCC3-Step, cells were collected and kept frozen until further used. All the purification steps were carried out at 0-4°C. The crude cell lysate was prepared from 6 g cell pellet (from 800 mL insect cell culture) by sonication in 50 mL T buffer (25 mM Tris-HCl, pH 7.5, 10% glycerol, 0.5 mM EDTA, 1 mM DTT, 0.05% IGEPAL, 1 mM PMSF and protease inhibitors) containing 300 mM KCl, 5 mM ATP, and 2 mM MgCl_2_. Lysates were clarified by centrifugation at 100,000 × g for 1 hr. The supernatant was incubated with 1 mL Ni-NTA affinity resin for 1 hr in the presence of 25 mM imidazole. The resin was sequentially washed with 100 mL T buffer containing 1 M KCl and then 50 mL T buffer containing 300 mM KCl, with all wash buffers containing 25 mM imidazole, 2 mM ATP and 2 mM MgCl_₂_. Bound proteins were eluted with 10 mL T buffer containing 200 mM imidazole, 300 mM KCl, 2 mM ATP, and 2 mM MgCl_₂_. The partially purified CX3 complex was passed through 1 mL HiTrap StepXT column, the column was further washed with 15 mL T buffer containing 300 mM KCl, 2 mM ATP, and 2 mM MgCl_₂_ (T300-ATP), and further eluted in same buffer also containing 50 mM biotin. Peak fractions were pooled and concentrated to 0.4 mL in an Amicon 30 concentrator and subjected to size exclusion chromatography in a Superdex200 Increase 10/300 GL column in (T300-ATP). Peak fractions containing CX3 complex were concentrated and snap frozen in liquid nitrogen and stored in −80°C.

#### DNA binding

For DNA binding, 2 nM of 5’ Cy5 labeled 80-nt ssDNA^15^ was incubated with the indicated concentration of purified BC and DX2 subcomplexes in 10 μL reaction buffer (50 mM Tris-HCl, pH 7.5, 120 mM KCl, 1 mM DTT, 1 mM ATP, 1 mM MgCl_2_ and 100 ng/μL BSA) for 10 min at 37°C. DNA-protein complexes were resolved on the 5% polyacrylamide gels in Tris-borate buffer (45 mM each) and gels were visualized using the ChemiDoc imaging system (Bio-Rad) and proportion of bound versus unbound DNA was quantified using the Fiji ImageJ software (v2.90/1.53t). The mean values were calculated from the data obtained from at least three independent experiments and plotted using the R (3.6.1) package ggpubr.

#### ATPase assays

The indicated combinations of BC and DX2 subcomplexes (0.5 μM) were incubated in the presence of ssDNA (20 μM nucleotides, phiX174 virion) in 10 μL of reaction buffer (20 mM HEPES, pH 7.5, 1 mM DTT, 1 mM MgCl_2_ and 30 mM KCl) containing 0.05 mM ATP with 0.25 μCi [γ-^32^P] ATP at 37°C. After 1 hr incubation, reaction was stopped by adding 10 μL of 0.5 M EDTA and the level of ATP hydrolysis was determined by layer chromatography on PEI cellulose F sheets (Millipore, 105579) in 375 mM potassium phosphate (pH 3.5), followed by phosphorimaging analysis. The mean values were calculated from the data obtained from at least three independent experiments and plotted using the R (3.6.1) package ggpubr.

#### Pull-down assay

Pulldown assays were performed by incubating 1 μg of the bait complex DX2 (RAD51D and XRCC2-FLAG) with 1 μg of either prey complex BC (RAD51B-His and RAD51C) or CX3 (RAD51C-His and XRCC3-Strep) in the presence of 2 μL α-FLAG resin (Pierce, A36804). Incubation was carried out in T buffer supplemented with 100 mM KCl, 2 mM ATP, and 2 mM MgCl_₂_ (T100-ATP) at 4°C for 1 hr with gentle agitation. After incubation, reactions were centrifuged to collect the flow-through (FT), and the resin was washed twice with T100-ATP. The resin bound proteins were analyzed by SDS-PAGE followed by Coomassie blue staining.

## Acknowledgements

This work was supported by the National Institutes of Health grants [R01 ES031796 (K.A.B.), R01 ES030335 (K.A.B.), GM115568 (S.K.O), GM128731 (S.K.O), R35 CA241801 (P.S.), P01 CA275717 (E.C.G. & P.S.), R01 CA293655 (S.K.O.), R35 GM153286 (J.O.K.), the American Cancer Society (DBG-1243615 to K.A.B), the Penn Center for Genome Integrity, the Basser Center for BRCA), the Department of Defense grants (BC201356 to K.A.B.). N.G. is supported by NIEHS (5T32ES019851-10). S.K.O. is the recipient of a CPRIT Rising Star Award (RR200030). S.L.H. is supported by an American Cancer Society Postdoctoral Fellowship.

## Supplemental Figures and Legends

**Supplementary Figure 1:**
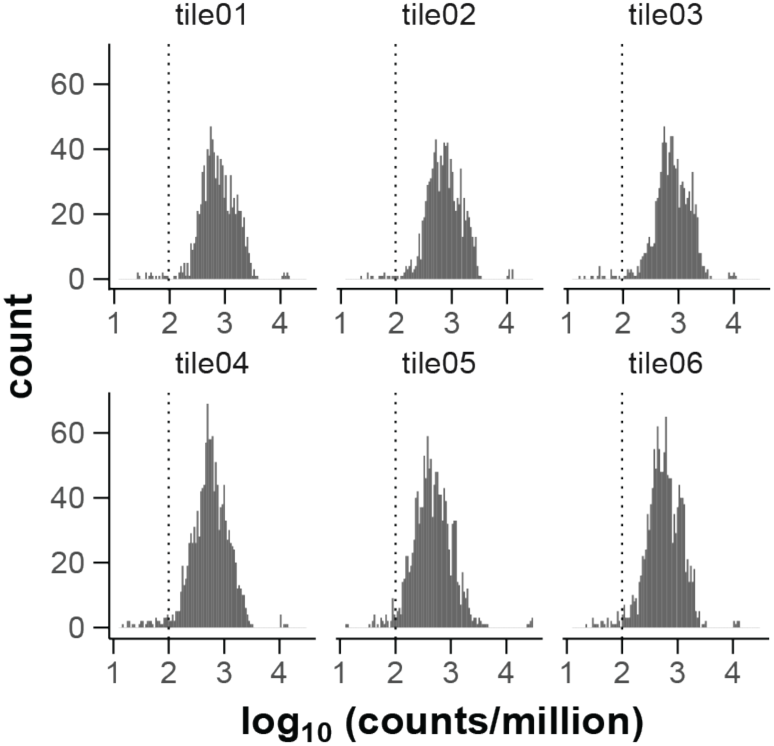
Mutant library uniformity. Histograms of *RAD51D* variant abundance, log10(counts/million counts), within pre-selection libraries for each mutagenesis tile. Cutoff line is at 100 cpm (1/10,000).

**Supplementary Figure 2:**
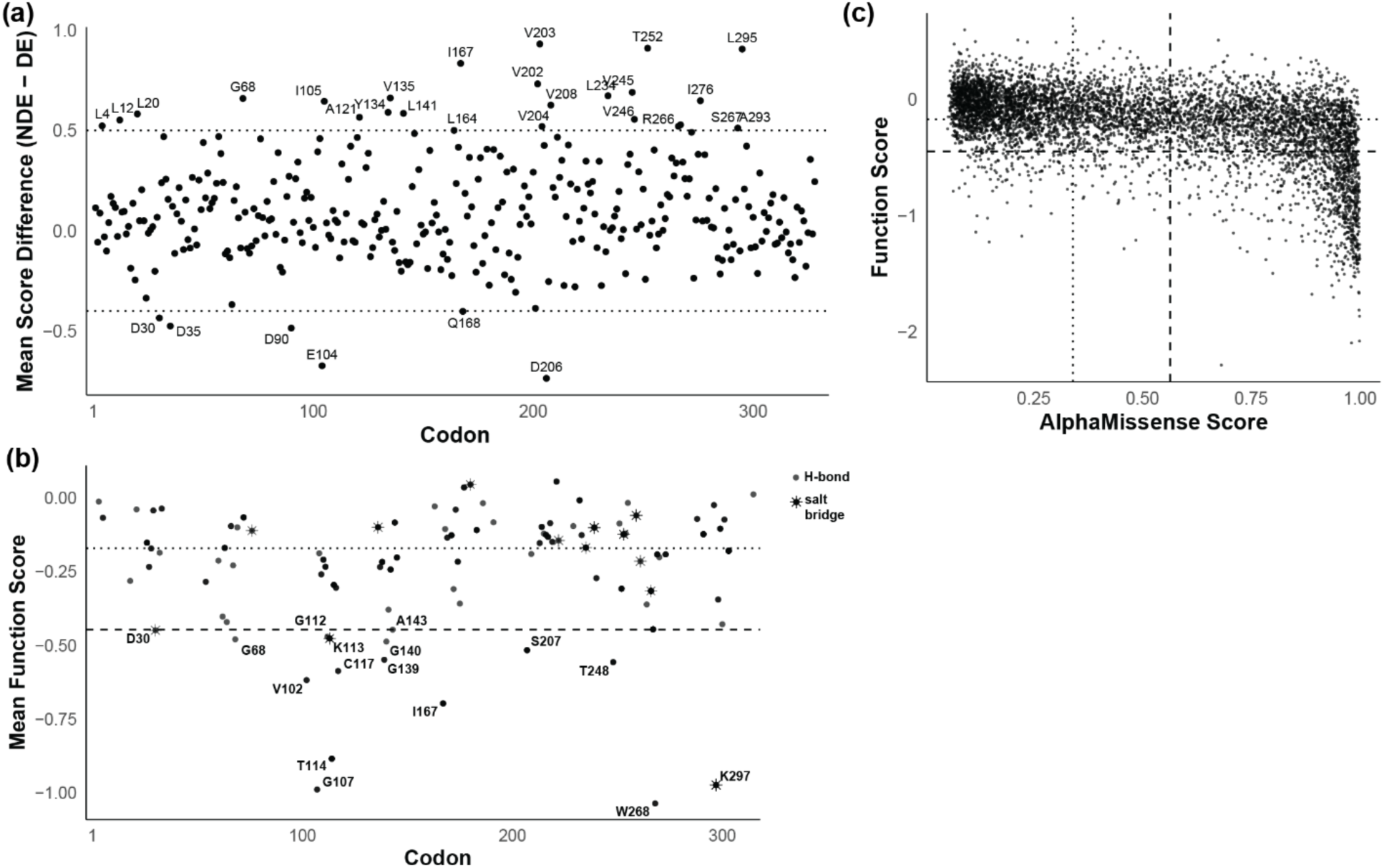
MAVE analysis of unique LOF residues. (**a**) Intolerance of negatively charged (i.e. Asp (D) and Glu (E)) residues throughout RAD51D, as measured by the difference between the mean function scores of D/E and non-D/E mutations. Residues with a difference ±0.5 are labeled. (**b**) Mean function scores of residues making contact with other proteins, ATP, or DNA via H-bonds or salt bridges (noted with circles or starbursts, respectively) in the BCDX2 complex (as predicted in PDB: 8GBJ). (**c**) AlphaMissense pathogenicity scores compared to MAVE function scores. AlphaMissense scores were binned as benign, pathogenic, and ambiguous using the published cutoff values. Counts of each plotted category are reported in the table.

**Supplementary Figure 3:**
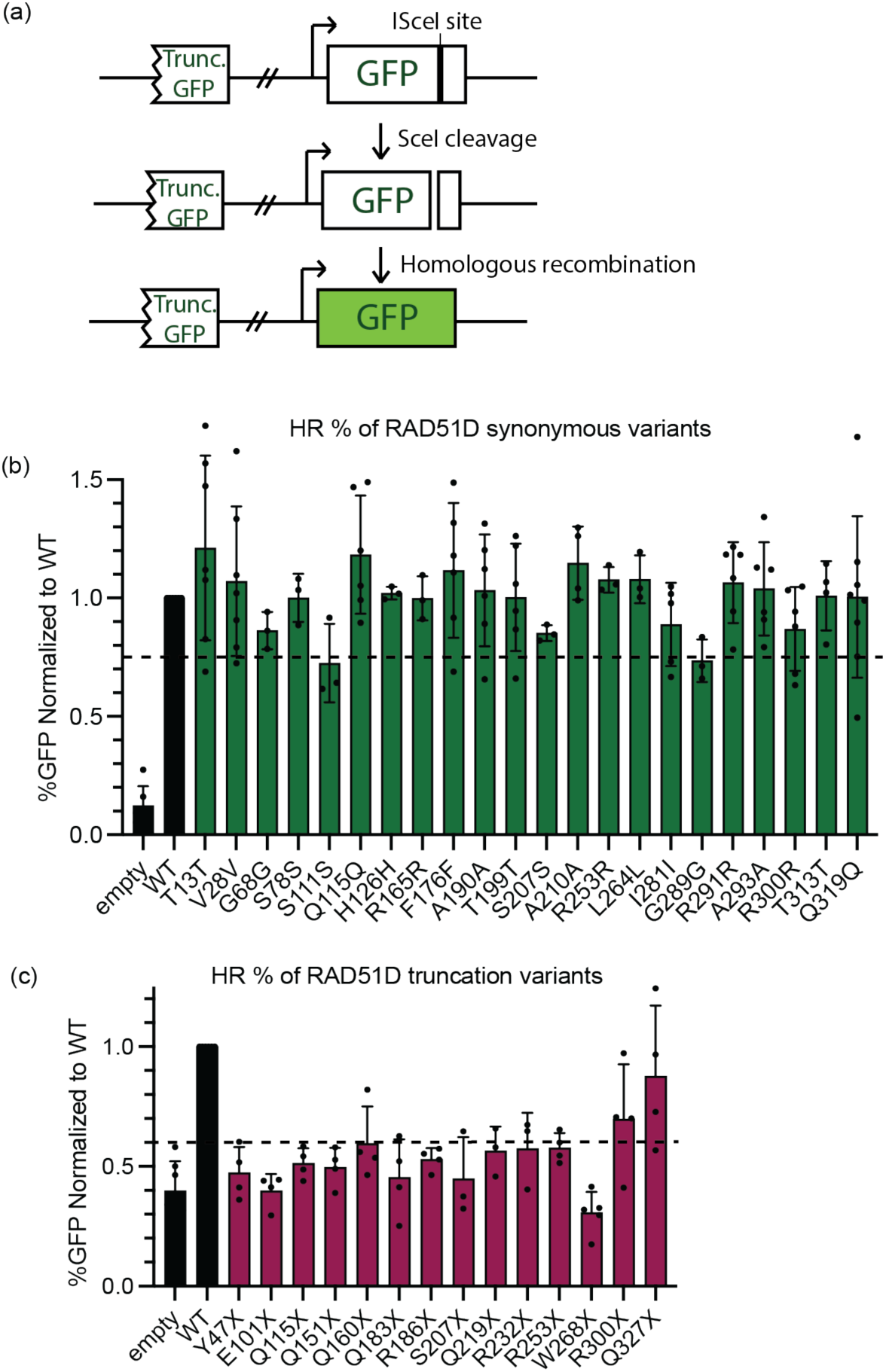
A series of synonymous and truncation variants were used to determine HR thresholding for benign and pathogenic variants. (**a**) Schematic of sister chromatid exchange assay to measure homologous recombination. In this assay, a cassette with a non-functional copy of GFP is integrated into RAD51D CRISPR/Cas9 U2OS cells. This GFP has a unique *I-SceI* restriction cut site. A DSB can be induced by expression of a plasmid expressing the I-SceI restriction enzyme. GFP expression is restored by use of a homologous template provided on the cassette following homologous recombination. (**b-c**) A plasmid with indicated synonymous or truncation variant was transiently transfected *RAD51D* CRISPR/Cas9 U2OS cells with a plasmid coding for the I-SceI restriction enzyme. The percentage of GFP+ cells was measured after three days, indicating a recombination event using a GFP fragment on the cassette. The HR proficiency threshold was determined based on comparison with the range of synonymous variants (green bars) to a wild-type *RAD51D* expressing plasmid (HR >0.75). The threshold for loss of HR function was calculated using the range of truncation variants compared to a wild-type *RAD51D* expressing plasmid, as <0.6 (indicated in red for the variants). Note that a subset of those variants analyzed here are replotted in Figure 5 as representative variants. The experiment was performed three to seven times with standard deviations plotted. An empty vector was used as a negative control.

**Supplementary Figure 4.**
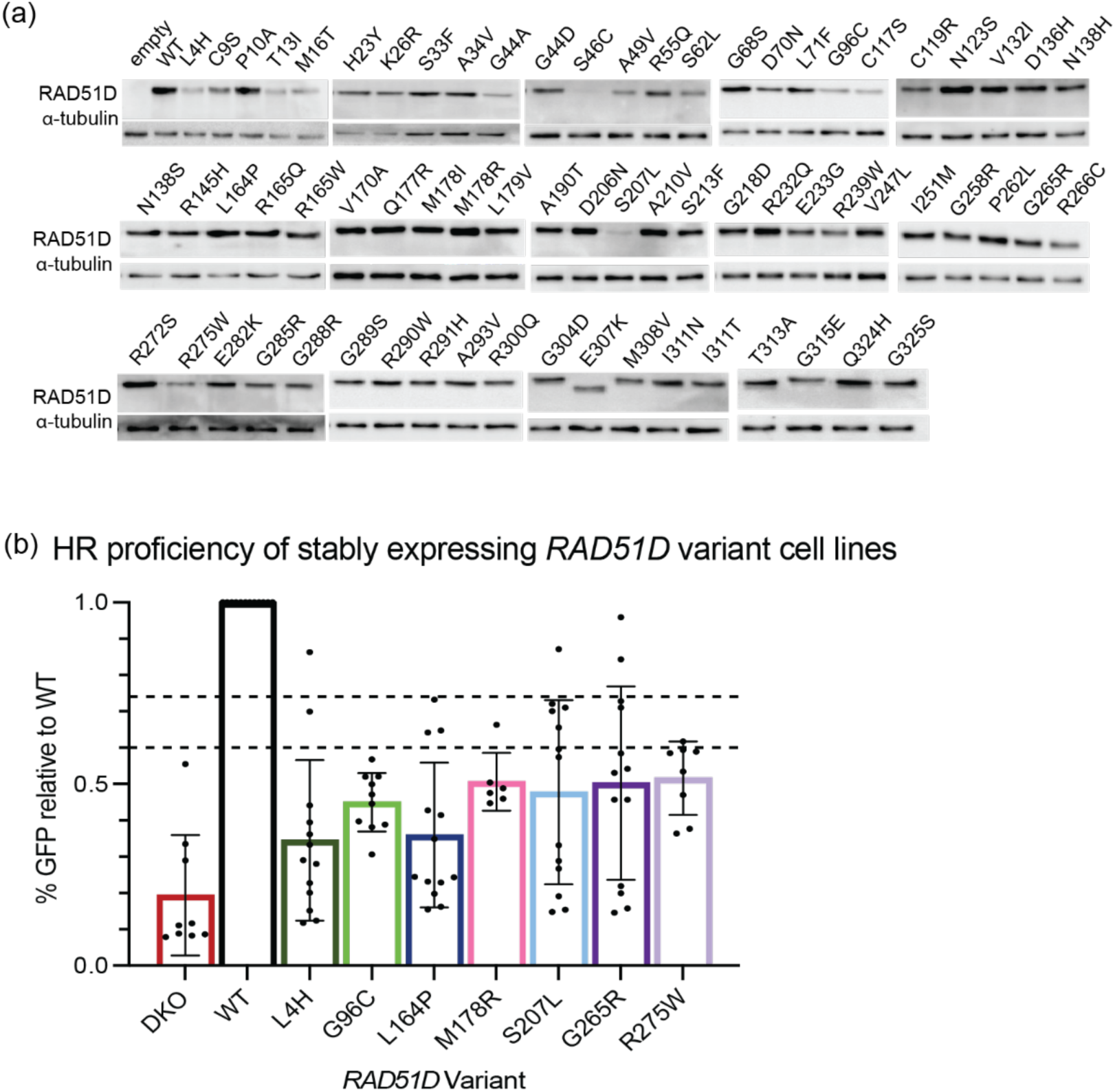
Expression and HR function of *RAD51D* variants in *RAD51D* CRISPR/Cas9 U2OS cells. (**a**) Western blot analysis of protein extract from U2OS cells expressing wild-type *RAD51D* or the indicated *RAD51D* variants was assessed using an anti-RAD51D antibody, and equal protein loading was assessed using an anti-tubilin antibody. Note that a subset of variants has reduced protein expression relative to WT RAD51D. Experiment performed in triplicate. (**b**) HR analysis of RAD51D variants that are stably expressed in the RAD51D KO cell line used for the clonogenic survival assays shown in Figure 4.

**Supplementary Figure 5.**
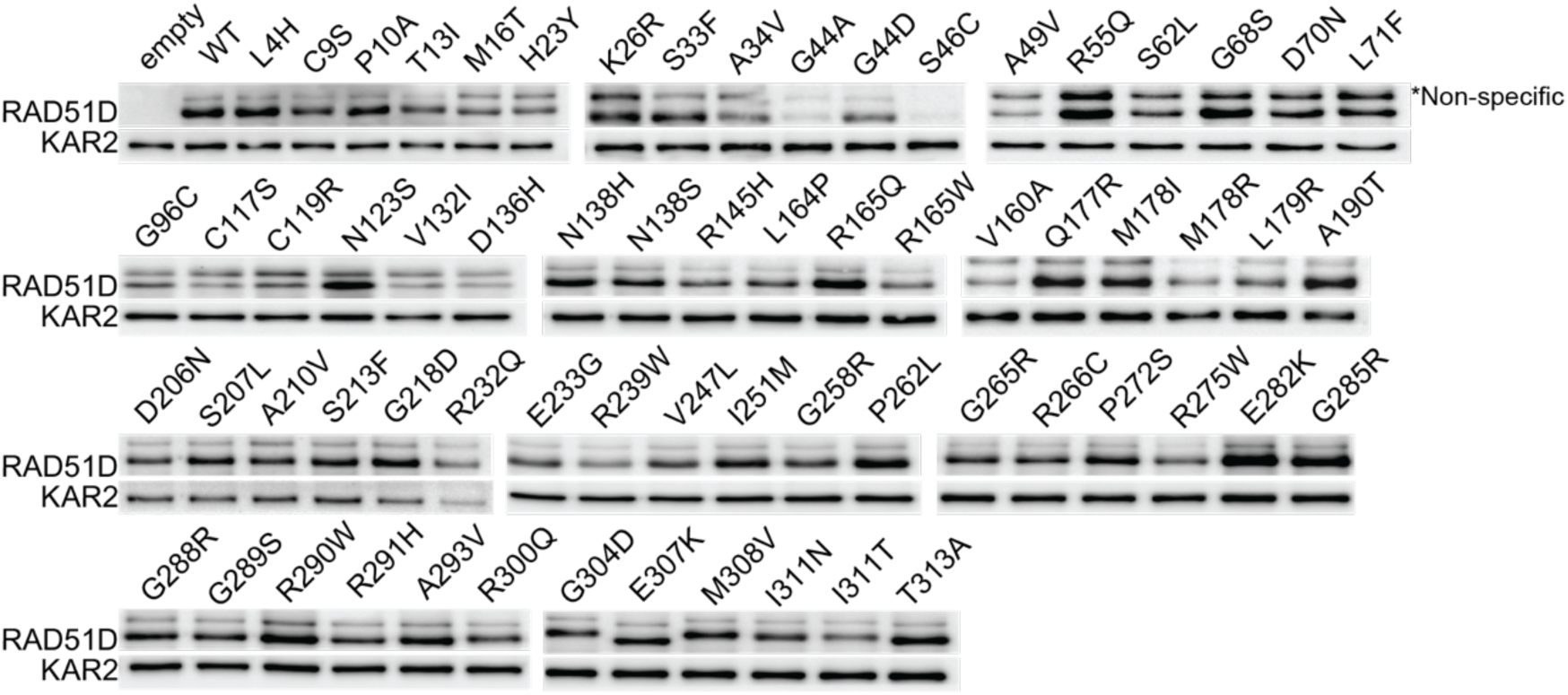
Western blot showing expression of RAD51D variants from the Y3H experiments. Western blot analysis of protein extract from yeast cells expressing wild-type *RAD51D* or the indicated *RAD51D* variants was assessed using an anti-RAD51D antibody, and equal protein loading was assessed using an anti-KAR2 antibody. Note that a subset of variants has reduced protein expression relative to WT RAD51D. Experiment performed in triplicate.

**Supplementary Figure 6.**
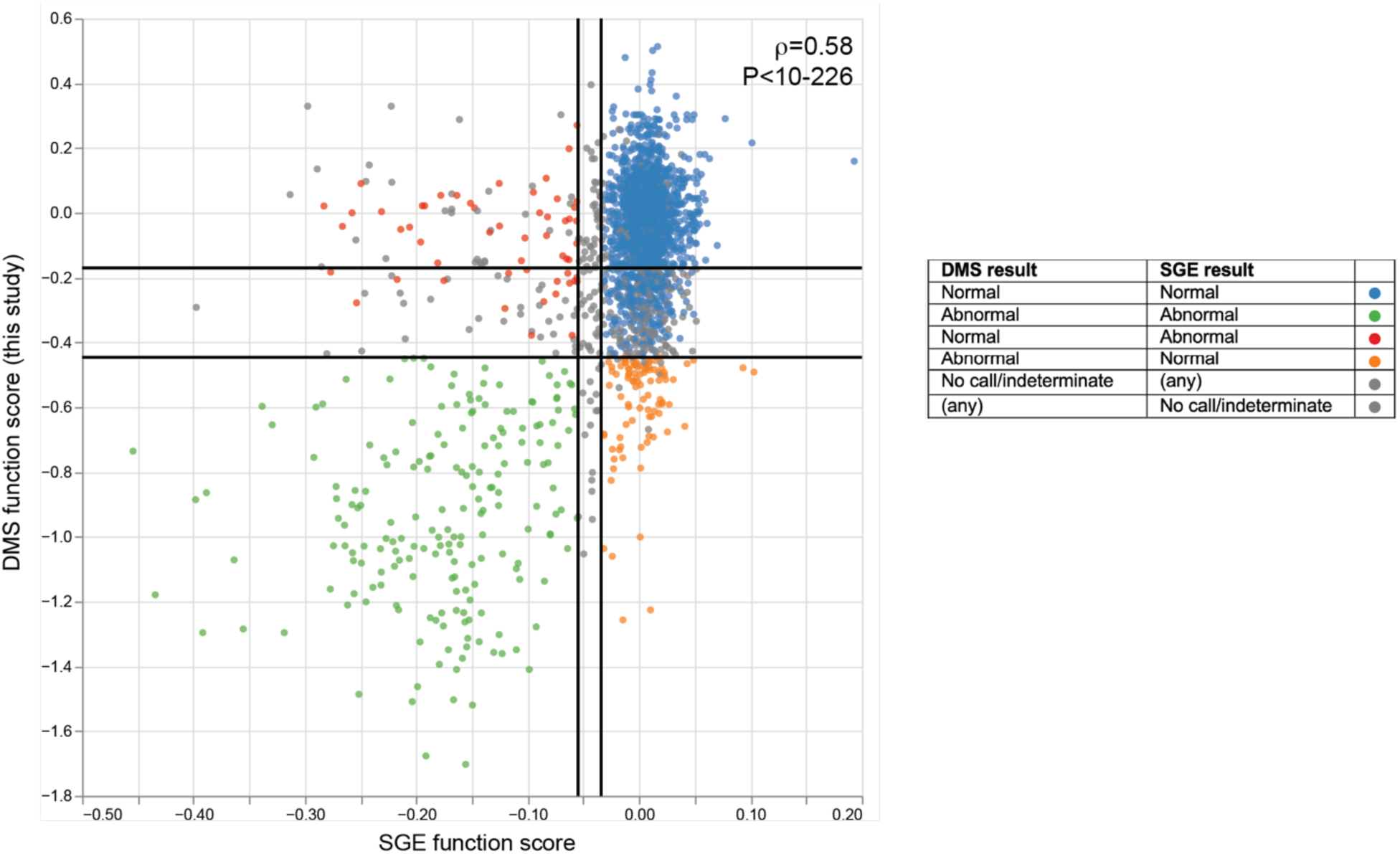
MAVE analysis from this study and Saturation Genome Editing (SGE) analysis on RAD51D are highly concordant. The MAVE (DMS) function score from this study was plotted relative to the SGE function score (*Casadei et al., in preparation*). Variants that both studies find to be normal are indicated in blue circles, variants that both studies find to be abnormal are indicated in green circles, conflicting results are shown in red or orange circles, and grey circles denote variants that received function scores from both studies but were deemed no-call and did not receive a classification from either one or both studies.

